# A clinically defined and xeno-free hydrogel system for regenerative medicine

**DOI:** 10.1101/2024.05.28.596179

**Authors:** John Ong, George Gibbons, Yee Siang Lim, Lei Zhou, Junzhe Zhao, Alexander W. Justin, Federico Cammarata, Ravisankar Rajarethinam, Colleen Limegrover, Sanjay Sinha, Andras Lakatos, Foad J. Rouhani, Yock Young Dan, Athina E. Markaki

## Abstract

Biofabricated scaffolds facilitate bona fide cellular interactions, cell type specification, and the formation of three-dimensional tissue architecture from human pluripotent stem cells (hPSCs). However, xenogenic biomaterials are poorly defined, and synthetic biomaterials remain underdeveloped and understudied, hindering regulatory approval for clinical use and preventing the translation of lab-grown therapies. Here, we describe a protein screen-based hydrogel system biofabricated from clinical-grade human components. We show that “Alphagel”, a base hydrogel comprising human embryonic matrices, supports the trilineage differentiation of hPSCs into neural, cardiac, and liver tissue. Alphagel is also shown to be biocompatible and biodegradable *in vivo*. Further, upon adding select proteins from maturing human foetal liver to Alphagel, we show that the resulting hydrogel (termed “Hepatogel”) enhances the differentiation of hPSC-derived hepatocytes (H-iHeps) compared with Matrigel. Importantly, when injected into mice livers, Hepatogel significantly improves the retention of H-iHeps compared to standard aqueous cell injections. Altogether, our results provide proof of concept that customisable and organ-specific hydrogel systems are a valuable tool for developing clinically translatable therapies for regenerative medicine and tissue engineering.

## 1. INTRODUCTION

Three-dimensional (3D) stem cell and organoid platforms recapitulate *in vivo* cellular and biomechanical interactions better than two-dimensional (2D) culture systems, enabling the development of novel diagnostics and therapeutics for disease (1,2). The spatial arrangement of stem cells and niche factors orchestrated by 3D biofabricated scaffolds promotes the self-organisation of cell masses and the formation of early organoids for tissue development. This geometry controls organoid patterning (3), and remains an essential factor that underpins our capability to engineer whole organs in vitro. Human pluripotent stem cells (hPSCs), which comprise induced pluripotent stem cells (iPSCs) and embryonic stem cells (ESCs), have the potential to generate any cell type of any organ. However, hPSC and organoid culture are heavily reliant on xenogenic matrices that often cause significant immune-related reactions when transplanted in humans, contributing to the failure of many tissue-engineered therapies in clinical trials (4).

Suitable alternatives to these 3D xenogenic matrices are lacking. Matrigel and Matrigel-derived matrices remain the predominant substrates used in the culture of liver cells and organoids from hPSCs (5–7). However, widely recognised issues are linked to these mouse sarcoma-derived matrices. Poor definability (8,9), batch-to-batch variability in growth factor concentrations (10,11), composition (12) and physical properties (11), immunogenicity (13), and pathogen transmission (14) are widely recognised issues that prevent clinical translation (15). As a result, synthetic biomaterials, such as polyethene glycol (PEG), polyacrylic acid (PAA), polycaprolactone (PCL), or polyacrylamide (PAAm), have been explored (16). However, synthetic components present specific challenges depending on the material used. For example, PEG-based biomaterials can exhibit rapid reaction kinetics, leading to inhomogeneous hydrogels (17). PAA hydrogels are mechanically weak, which limits their use as scaffolds in tissue engineering (18). PCL hydrogels exhibit poor cell adhesion and involve toxic reagents during synthesis (19). PAAm hydrogels require complex chemical or photo-crosslinking (20), precluding gelation and retention at target sites for convenient and effective cell delivery (8,21). Without biomodifications, synthetic biomaterials lack the biochemical cues required for efficient hPSC culture and differentiation (16,22). Given these challenges, synthetic biomaterials have rarely been used to bioengineer liver cells and tissue from hPSCs (5).

Recently, attempts have been made to deconstruct components of Matrigel and conjugate these with various polymeric backbones (23,24) for liver bioengineering. However, these are unsuitable for hPSC culture because some components can adversely affect stem cell pluripotency by causing predilection or spontaneous differentiation towards particular lineages (25–27). For example, Laminin 111, a major component of Matrigel, can promote epithelial-to-mesenchymal transition in human ESCs when its β1–LN–LE1-4 fragment triggers the α3β1-integrin/extracellular matrix metalloproteinase inducer complex (26).

At present, there are no clinical-grade ECM substrates that support the 3D culture of hPSCs and their differentiation into liver tissue (5). Thus, we biofabricated a xenofree, clinically compatible system that mimics the stages of embryonic development and liver organogenesis. We hypothesised that hPSC-derived target cells would more closely resemble primary cells when cultured in customised organ-specific hydrogels rather than generic biomaterials such as Matrigel or synthetic hydrogels. In that vein, we aimed to biofabricate a liver extracellular matrix (ECM)-specific hydrogel as a proof-of-concept. Taking a reductionist approach, we performed a focused protein screen and created a base hydrogel (termed “Alphagel”) that supported hPSC culture and trilineage differentiation in 3D. We then added selected proteins from maturing liver ECM to Alphagel. The resulting hydrogel, termed “Hepatogel”, produced hPSC-derived hepatocytes that were closer to maturing human hepatocytes than Matrigel, and it improved the engraftment of hPSC-derived hepatocytes when injected into the livers of immunocompromised mice. In so doing, a clinically defined (composition suitable for clinical use) hydrogel system was achieved. A summary of the workflow is illustrated in Fig.1(A). Here, we provide evidence that organ-specific hydrogels yield better end-target cells than generic hydrogels and that this system is a useful tool for developing organ-specific, effective therapies for regenerative medicine.

**Figure 1.**
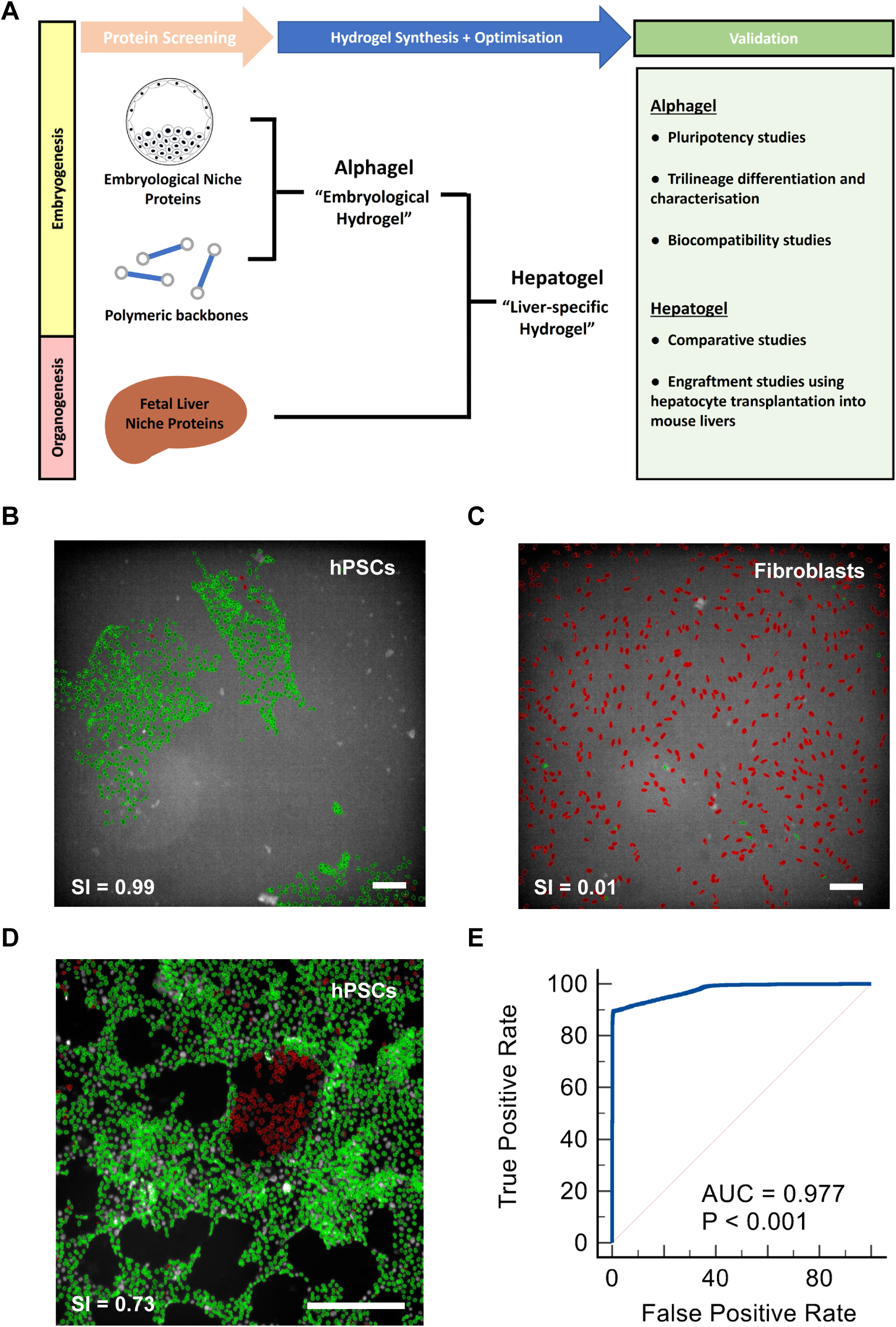
A) Schematic of workflow. B) Automated pipeline identifying hPSC colonies as pluripotent stem cells; green = positive selection by machine learning, scale bar = 100 μm. C) Machine learning identifying human fibroblasts as non-pluripotent cells (red); scale bar = 100 μm. D) Machine learning discriminating undifferentiated hPSCs (green) from spontaneously differentiated hPSCs (red) after intentionally extended culture to trigger spontaneous differentiation (Day 9), scale bar = 200 μm. E) A Receiver Operator Characteristic curve showing excellent discriminatory performance of our machine-learning platform with an Area Under the Curve (AUC) of 0.977.

## 2. MATERIALS AND METHODS

### 2.1 Focus Protein Screening

Protein screening was performed using the Opera Phenix High-Content High-Throughput (HCHT) screening system (PerkinElmer, USA). Nuclear morphology was measured using DAPI, and the mean fluorescent intensity (MFI) of Octamer-binding transcription factor 4 (OCT4), homeobox transcription factor Nanog (NANOG), and sex determining region Y-box 2 (SOX2) were used as stem cell pluripotency markers. Machine learning thresholds were established using the MFI of OCT4, NANOG, and SOX2 in hPSCs (positive control) cultured on Geltrex (Thermo Fisher, #A1413302) and human fibroblasts (negative control). hPSCs were then plated on 96-well tissue culture-treated plates (Greiner Cellstar, #M0812) coated with varying substrates at 4 different concentrations. Our pipeline first identified single cells based on nuclear properties, then selected hPSCs based on the MFI of OCT4, NANOG, and SOX2. A brief outline is described in Supplementary Figure 1. The measure of pluripotency within a stem cell population was then defined as the “Stemness Index” (SI), which was calculated by:

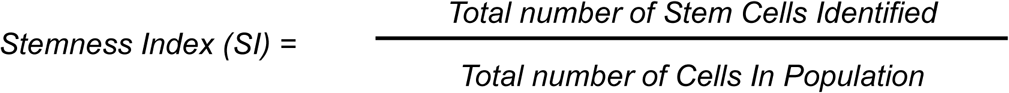

The SI values range from 0 to 1, and the equation was incorporated into the pipeline. Positive hits were checked using brightfield microscopy.

### 2.2 Two-dimensional (2D) cell culture

Human ESCs (KCL033) and human iPSCs (RMA and BOB cell lines) gifted by Dr Tamir Rashid (King’s College London) were used for the focused protein screening. Commercial ESC cell lines (H1 and H9, WiCell #WA01 and #WA09, respectively) and 1 episomal iPSC cell line (Thermo Fisher, #A18945) were used for validation. hPSCs were maintained on Geltrex-coated plates in feeder-free mTeSR-1^TM^ and mTeSR Plus^TM^ media (Stemcell Technologies #85850 and #100-0276) using an established protocol (28). During maintenance, cells were split at 70-80% confluence using Accutase (Millipore, #SCR005), washed with DMEM/F12 (Life Technologies, #11330032) and re-seeded onto Geltrex-coated 6-well plates (Greiner, #657160) at a ratio of 1:6 to 1:8. 1mM Y27632 ROCK inhibitor (Stratech Scientific, #S1049-SEL-2mg) was used to increase cell attachment. Media was supplemented with penicillin and streptomycin (Millipore, #TMS-AB2-C). Trypan Blue (Sigma, #15250061) was used to assess cell viability.

Human dermal fibroblasts (Sigma, #106-05a) were cultured in uncoated tissue culture flasks (Corning, #CLS3275) or in Geltrex-coated 6-well plates. Fibroblasts were maintained in DMEM/F12 media (Life Technologies, #11330032), 10% foetal bovine serum (Sigma, #A9418), and 1% penicillin and streptomycin. Fibroblasts were passaged using Trypsin (Thermo Fisher Scientific, #11538876).

Primary adult human hepatocytes (PHH) were obtained commercially (Lonza, #HUCPI) and used as positive controls of hepatocyte function. These were ethically sourced, quality-controlled, characterised and came from three 3 different non-diseased donors; Lot no. HUM182701, HUM182641, and HUM191381. PHH were thawed, plated on collagen-1-coated plates, and recovered for at least 24 hr in Hepatocyte Culture MediaTM (HCM), including the complete HCM bullet kit (Lonza, #CC-3198), according to the supplier’s protocol. Experiments commenced after 24 – 48 hr of cell recovery.

### 2.3 Three-dimensional (3D) cell culture

H1, H9, and Episomal cell lines were cultured in 3D domes of Matrigel, fibrin-only gels, or Laminin 521-enriched fibrin hydrogels (FLGs), including Alphagel. Domes were plated on hydrophobic non-tissue culture-treated hydrophobic plates under normoxic conditions at 37 °C. 50 μl domes were used for 24-well plates (Sarstedt, #83.3922.500) or 25 μl domes for 96-well plates (Ibidi, #89621). Feeder-free mTeSR-1^TM^ and mTeSR Plus^TM^ media (Stemcell Technologies #85850 and #100-0276) were used for routine stem cell maintenance. hPSCs designated for animal experiments were maintained in xeno-free TeSR^TM^ 2 media (Stemcell Technologies, #05860). Cells were split every 5-6 days using TrypLE Express (Gibco, #12604013). Each dome required 20-25 min in TrypLE Express with manual pipetting for successful dissociation. Domes were then reloaded with 12,500-22,500 cells, depending on the experiment.

For experiments investigating stem cell culture media on hPSC pluripotency in Alphagel, TeSR-E8^TM^ culture media (Stemcell Technologies, #05990), E8^TM^ culture medium (Thermo Fisher, #A1517001), and TeSR-E8TM culture medium (Stemcell Technologies, #05990) were used in addition to those described above.

### 2.4 Cardiac Differentiation in 3D

hPSCs were transitioned to E8^TM^ culture medium (Thermo Fisher, #A1517001) supplemented with 2.25 ng/ml of Fibroblast Growth Factor-2 (FGF-2: Qkine, #Qk053), and 1.74 ng/ml of Transforming Growth Factor Beta (TGF-β: Bio-techne, #7754-BH) as previously described (29). When hPSC colonies reached the desired number and size, cardiac differentiation was initiated by replacing the FGF-2 and TGF-β enriched E8 maintenance media with CDM-BSA (Sanjay Sinha Lab, University of Cambridge) containing 20 ng/ml of FGF-2, 10 μM of 2-(4-Morpholinyl)-8-phenyl-4 H-1-benzopyran-4-one (Ly294002: Promega, #V120A), 50 ng/ml of Activin A (Stemcell Technologies, #78132.2), and 10 ng/ml of Bone Morphogenic Protein-4 (BMP-4: RnD system, #314-BP-050) (30). After 42 hr (Day 1.5), this was replaced with CDM-BSA containing 8 ng/ml of FGF-2, 10 ng/ml of BMP-4, 1μM of retinoic acid (Sigma, #R2625) and Wnt/β-catenin signalling inhibitor endo-IWR1 (R&D systems, #3532). Media was refreshed after 48 hr (Day 3.5). After another 48 hr (Day 5.5), the media was replaced with CDM-BSA containing 8 ng/ml of FGF-2 and 10 ng/ml of BMP-4. Standard CDM-BSA media was then refreshed every 48 hr (Day 7.5 onwards).

### 2.5 Neuronal Differentiation in 3D

hPSCs were transitioned into Stem Flex media (ThermoFisher, # A3349401) with 10μM Y-27632 (Tocris, #1254) before initiating neuronal differentiation. The media was refreshed daily. When pluripotent spheroids were of sufficient number and size in Alphagel, the media were switched to N2B27 media (in-house) supplemented with 5 μM SB-431542 (Tocris, #1614), 200 nM LDN-193189 (Miltenyi, #130-103-925), and 2 μM XAV-939 (Tocris, #3748) (31). Media with these factors was refreshed daily for ten days. After this, cells were maintained in N2B27 media for an additional 6 days.

### 2.6 Hepatic Differentiation in 3D

Hepatocyte differentiation was performed using an established protocol (28). Briefly, hPSCs were cultured in RPMI (Thermo Fisher, #22400089) supplemented with 2% B27 minus insulin (Thermo Fisher, #A1895601), 100 ng/ml of Activin A, 20 ng/ml of FGF-2 (Thermo Fisher, #PHG0023) and 10 ng/ml of Bone Morphogenic Protein-4 (BMP-4: R&D system, #314-BP-050) to initiate Day 1 and 2 of differentiation. From differentiation days 3-5, RPMI supplemented with 2% B27 minus insulin and 100 ng/ml Activin A was used. Hepatic endoderm specification was initiated on differentiation days 6-10 using RPMI supplemented with 2% B27, insulin (Thermo Fisher, #17504044), 10 ng/ml FGF-2, and 20 ng/ml BMP-4, followed by RPMI with 2% B27, insulin, and 20 ng/ml hepatocyte growth factor (HGF: Peprotech, #100-39-250) for differentiation days 11-15. Maturation of hepatic endoderm was initiated on differentiation days 16-21 using complete Hepatocyte Culture Media (HCM) supplemented with 10 ng/ml Oncostatin M (OSM: R&D systems, # 8475-OM-050), 20 ng/ml HGF and 0.1 μM Dexamethasone (Thermo Fisher, #A17590.06).

### 2.7 Synthesis of fibrin hydrogels

Human fibrinogen from non-diseased donors (Merck Millipore, #341576-1GM) was reconstituted in sterile 0.9% saline to a 67 mg/mL stock concentration. Human thrombin (Sigma, #T6884-100UN) was reconstituted in sterile 0.9% saline supplemented with human serum albumin (0.1% w/v), giving a stock concentration of 50 U/ml. DMEM/F12 was then added to varying volumes of fibrinogen and thrombin to achieve the desired final fibrinogen concentration in the fibrin gels. Final fibrinogen concentrations in the hydrogels we tested ranged from 1 mg/ml to 10 mg/ml. The final thrombin concentration in these hydrogels was maintained at 1 U/ml. For cell experiments, hPSCs were loaded into the DMEM/F12 compartment.

### 2.8 Synthesis of Alphagel and Hepatogel

Alphagel and Hepatogel were synthesised using similar methods of fibrin-laminin enrichment previously reported (23,32) with adjustments. Briefly, for acellular Alphagel domes, human recombinant laminin 521 (Biolamina, #CT521-0501/LN521-05, 0.1 mg/mL), human fibrinogen and DMEM/F12 were incubated in sterile Eppendorfs (Elkay Lab, #MICR-050) at 37 °C for 15 min. The solution was then mixed and incubated for a further 15 min. Lastly, thrombin diluted in DMEM/F12 at room temperature was added for gelation. The final composition of Alphagel was a 50:50 v/v mixture of 0.05 mg/mL Laminin 521 and 2.5 mg/mL fibrinogen (final concentration), crosslinked with 1 U/ml thrombin.

For Hepatogel, Laminin 411 (Biolamina, #LN411, 0.1 mg/mL) and Laminin 111 (Biolamina, #LN111, 0.1 mg/mL) were purified using Amicon Ultra-0.5 centrifugal filter units (Merck Millipore, UFC5003) and reconstituted in sterile phosphate-buffered saline (Sigma, #806552). Laminin 521, Laminin 411, and Laminin 111 (all at 0.1 mg/mL stock concentrations) were mixed at a 5:1:2 ratio (v/v/v). The laminin mixture was subsequently combined 4:1 with fibrin precursor solution (2.5 mg/mL fibrinogen and 1 U/mL thrombin), as described above, yielding a final total laminin concentration of 0.08 mg/mL within the gel. This corresponded to final concentrations of 0.05 mg/mL Laminin 521, 0.01 mg/mL Laminin 411, and 0.02 mg/mL Laminin 111.

For cell experiments, hPSCs were loaded in the laminin(s)-DMEM/F12-fibrinogen compartment and incubated as described above. The thrombin-DMEM/12 compartment was added for polymerisation. The cell-laden mixture was seeded as domes on hydrophobic plates and kept inverted for 30 min to allow cell suspension and gelation to complete fully. Thereafter, plates were turned over, and media was added. The media was refreshed at a frequency determined by experiments.

### 2.9 Scanning Electron Microscopy (SEM)

Hydrogels were mounted and gelled in 444 ferritic stainless-steel scaffolds (Nikko Techno Ltd., Japan) for mechanical support. An hour after gelation, samples were fixed in 10% formalin (VWR International, #11699455) and left in a fume hood overnight. Samples were then washed thrice with PBS and dehydrated in increasing concentrations of ethanol-de-ionised (DI) water mixtures: 30, 50, 70, 80, 90, 95 and 100% ethanol (33). Samples were submerged in each ethanol-DI mixture for 30 min. The final step with 100% ethanol was repeated twice. For dehydration, samples were immersed in increasing concentrations of hexamethyldisilazane (HMDS) in ethanol mixtures in a fume hood: 33.3%, 66.6%, and 100% HMDS (Sigma, #440191), respectively. Samples were immersed in each HMDS-ethanol mixture for 40 min. The final step involving 100% HMDS was repeated twice, and then the samples were left submerged in 100% HMDS overnight in a fume hood until completely dry the next day.

Dried samples were gold-plated before imaging with the Evo LS 15 (Zeiss, Germany) scanning electron microscope.

### 2.10 Immunofluorescence (IF) staining

Monolayer cells were washed twice with PBS (100 μl per well in 96-well plates) and fixed with 4% paraformaldehyde (PFA, Insight Biotechnology, #sc-281692) for 20 min at 4 °C. After two further PBS washes. 50 μl of blocking solution containing 3% donkey serum (Sigma, #D9663-10ML), 0.1% Triton-X (Sigma, #X100-500ML), 1% BSA (Sigma, #12657), and 95.9% PBS was added to each well. After 30 min at room temperature, this was aspirated, and the primary antibodies were added without washing. The list of primary antibodies is listed in Table 1. After 2 hr, the primary antibodies were aspirated, and two washes were performed (100 μl/well PBS per wash). 50 μl/well of secondary antibodies were added, and the samples were incubated at room temperature in the dark for 2 hr. The list of secondary antibodies is listed in Table 2. After 2 hr, secondary antibodies were aspirated, and cells were washed twice with PBS (100 μl/well). DAPI solution (NucBlue^TM^ Fixed, Life Technologies #R37606) was then added, and imaging was performed within 24 hr.

**Table 1:**
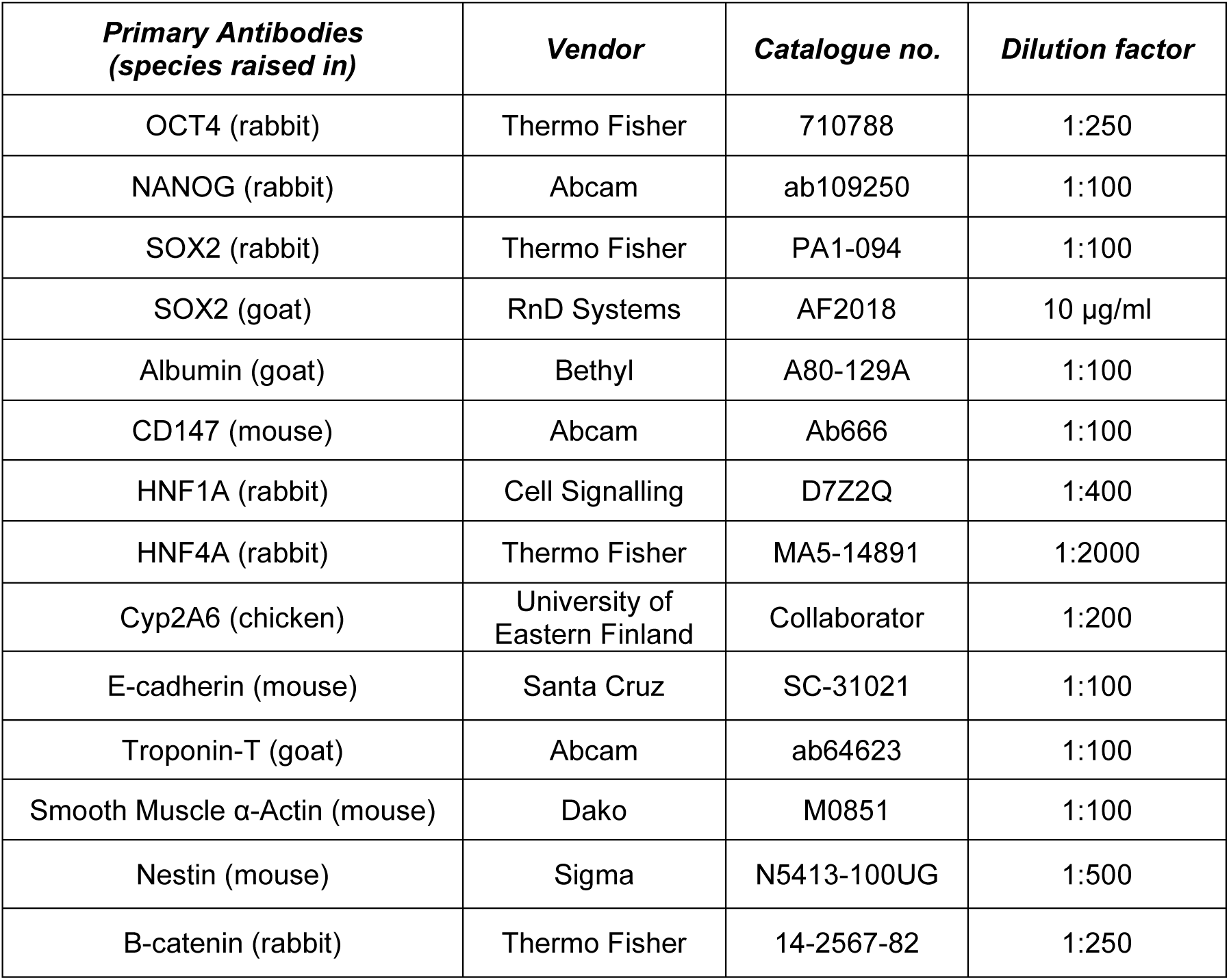
List of primary antibodies.

**Table 2:**
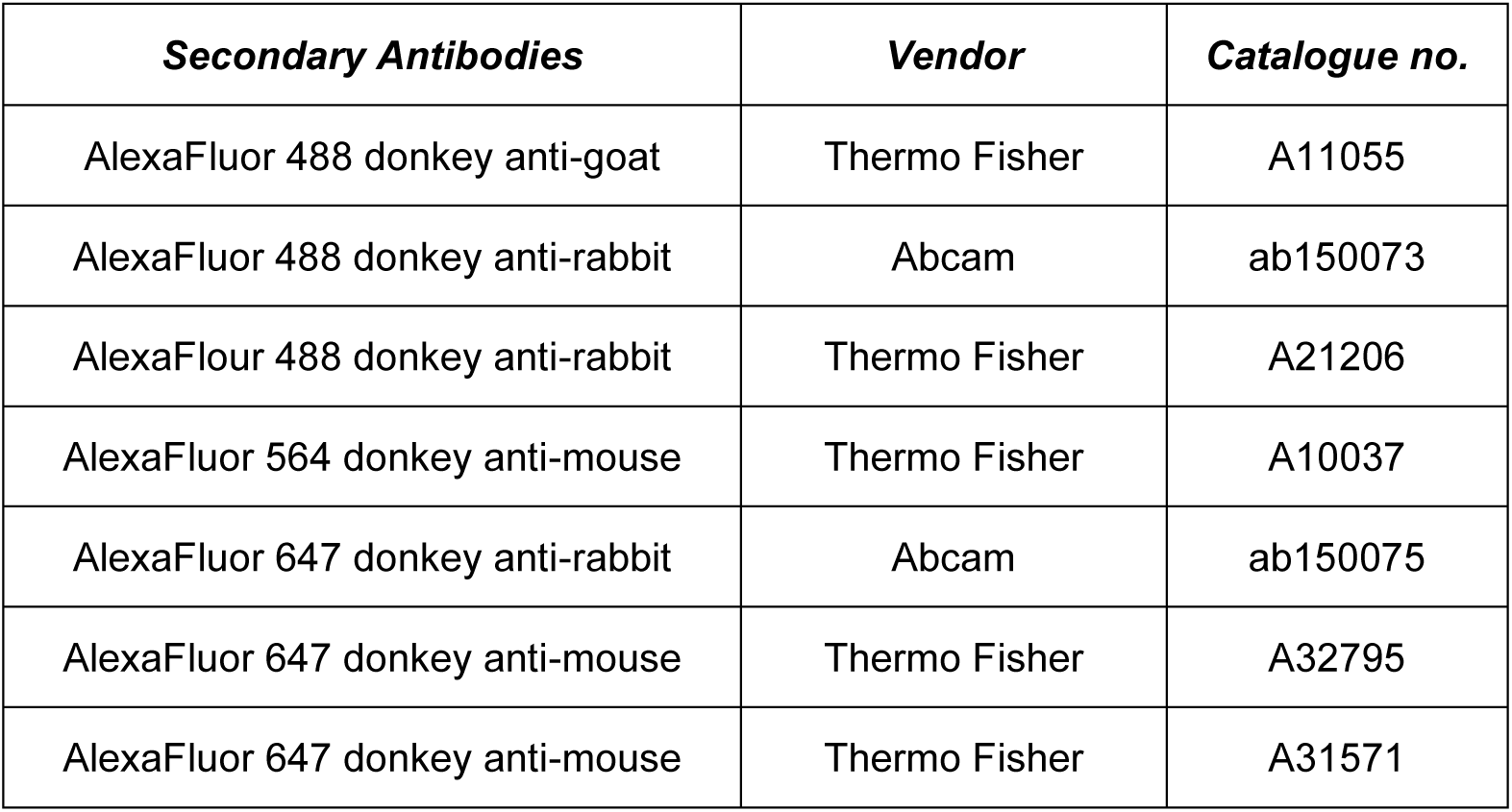
List of secondary antibodies.

For 3D cardiac and hepatic experiments, cell-laden hydrogel domes underwent two 15-minute washes with PBS (1 ml/well in a 24-well plate). Samples were then fixed for 60 min at 4°C with 4% paraformaldehyde (1 ml/well), followed by three 15-min washes with PBS (1 ml/well). Cells were permeabilised and blocked in 1% (w/v) bovine serum, 3% donkey serum, and 0.1% Triton in sterile PBS for one hour at room temperature. The blocking solution was aspirated, and primary antibodies (1ml/well) were added without washing. Samples were incubated overnight at 4 °C. Primary antibodies were aspirated the following day, and samples underwent two fifteen-minute PBS washes. Secondary antibodies were added, and samples were incubated in the dark at room temperature for ≥3 hr before aspirating. DAPI solution was then added, and images were acquired at least 30 min after its addition. For 3D neural experiments, hydrogels were fixed in 4% PFA for 45 min at room temperature, then transferred to a 30% w/v sucrose solution (Sigma-Aldrich, #S0389) and incubated for 24 hr. Sucrose-saturated samples were placed in Optimal Cutting Temperature (OCT) medium (Thermo Fisher, #23-730-571) and stored at-20 °C. Frozen blocks were cut into 12 μm sections and mounted on coated slides (Thermo Fisher, #J1800AMNZ). Sections were then blocked and permeabilised in a 0.3% Triton solution containing 10% normal goat serum at room temperature for an hour. Primary antibodies diluted in 0.1% Triton containing 5% normal goat serum were then added and incubated overnight at 4 °C. The following day, sections were washed with PBS, and secondary antibodies were added and incubated for 1 hour at room temperature before aspirating. Samples were then incubated in DAPI solution for 15 min and then mounted using FluorSave (VWR, #345789).

For IF staining of tissue sections, glass slides with paraffin-embedded sections were rehydrated in xylene (5 min x3) and transferred to reducing concentrations of ethanol (100% for 5 min x2, 90% for 2 min, and 70% for 2 min). Thereafter, slides were washed in distilled water for 5 min, transferred to pH 6.0 antigen retrieval buffer, and heated to 121 °C for 15 min. The slides were re-washed with distilled water after cooling. TrueBlack® (Biotium, #23007) dissolved in 70% ethanol (1X) was added to the samples for 30 sec to reduce autofluorescence. Slides were then rinsed with distilled water, and the blocking solution was added. Samples were left for 30 min at room temperature then the blocking solution was discarded. Without washing, goat anti-human albumin primary antibody dissolved in sterile PBS to a final concentration of 1:100 (as described above) was added and slides were incubated at 4 °C overnight. The next day, the slides were washed with distilled water, and 1:200 Alexa Fluor 647 donkey anti-goat secondary antibody was added. Slides were then incubated in the dark at room temperature for 1 hour, then washed again before adding Fluoroshield^TM^ mounting media containing DAPI (Sigma, #F6057).

### 2.11 Image Acquisition

2D IF and brightfield images of hPSCs were acquired using the Opera Phenix HCHT screening system (PerkinElmer, USA). An exposure time of 100ms with 90% excitation was kept constant during screening. 3D images of cardiac and hepatic cells and tissue were captured using an LSM710 (Zeiss, Germany) laser-scanning confocal microscope at x10 and x20 objectives with the corresponding filters. Bright-field images were acquired using an Axio-Obsorber Z1 (Zeiss, Germany) fluorescence microscope. 5-(and-6)-Carboxy-2’,7’-Dichlorofluorescein Diacetate (CDFDA) uptake and excretion images in hPSC-derived hepatocytes (iHeps) were acquired using an SP-DM8 (Leica, Germany) microscope. 3D images of neuronal cells were acquired using a DM 6000 B (Leica, Germany) microscope.

### 2.12 Young’s modulus measurements

Fibrin-only and FLGs were examined under compression using a customised “see-saw” method that was previously described (34), and loads were applied by pumping liquid via a syringe pump. Briefly, two arms of a see-saw were first balanced before counterweights were added onto the free-extending arm to match the height of the loading platen with the sample height. Next, the load was ramped up at a constant rate of 10^-3^ N s^-1^ by pumping liquid via a PHD ULTRA syringe pump (Harvard Apparatus, USA) into a container mounted on the loading platen. The resulting displacement was measured using a biaxial laser micrometre (resolution of ±3 µm, Keyence, #TM-040). Nominal stresses (σ) and strains (ε) were plotted, and a line was least-squares fitted to the data points. The through-thickness Young’s modulus was calculated from the tangent slope of the stress-strain curves (n=5 in each group).

### 2.13 Gene expression analyses by quantitative polymerase chain reaction (qPCR)

Ribonucleic acid (RNA) was extracted using TriZOL^TM^ (Thermo Fisher, #15596026) and purified using the Direct-zol RNA Miniprep Kit (Zymo Research, #R2050-R2053). cDNA was synthesised using Superscript Reverse Transcriptase IV (Thermo Fisher, #18090010) according to the manufacturer’s protocol. Quantitative PCR was performed on a QuantStudio 5 (Thermo Fisher, #A34322I) using the KAPA SYBR FAST qPCR kit (Roche, #KK4601). Data were analysed using the 2^(ΔΔCt) method. Sequences of the primer pairs are listed in Table 3.

**Table 5.3:**
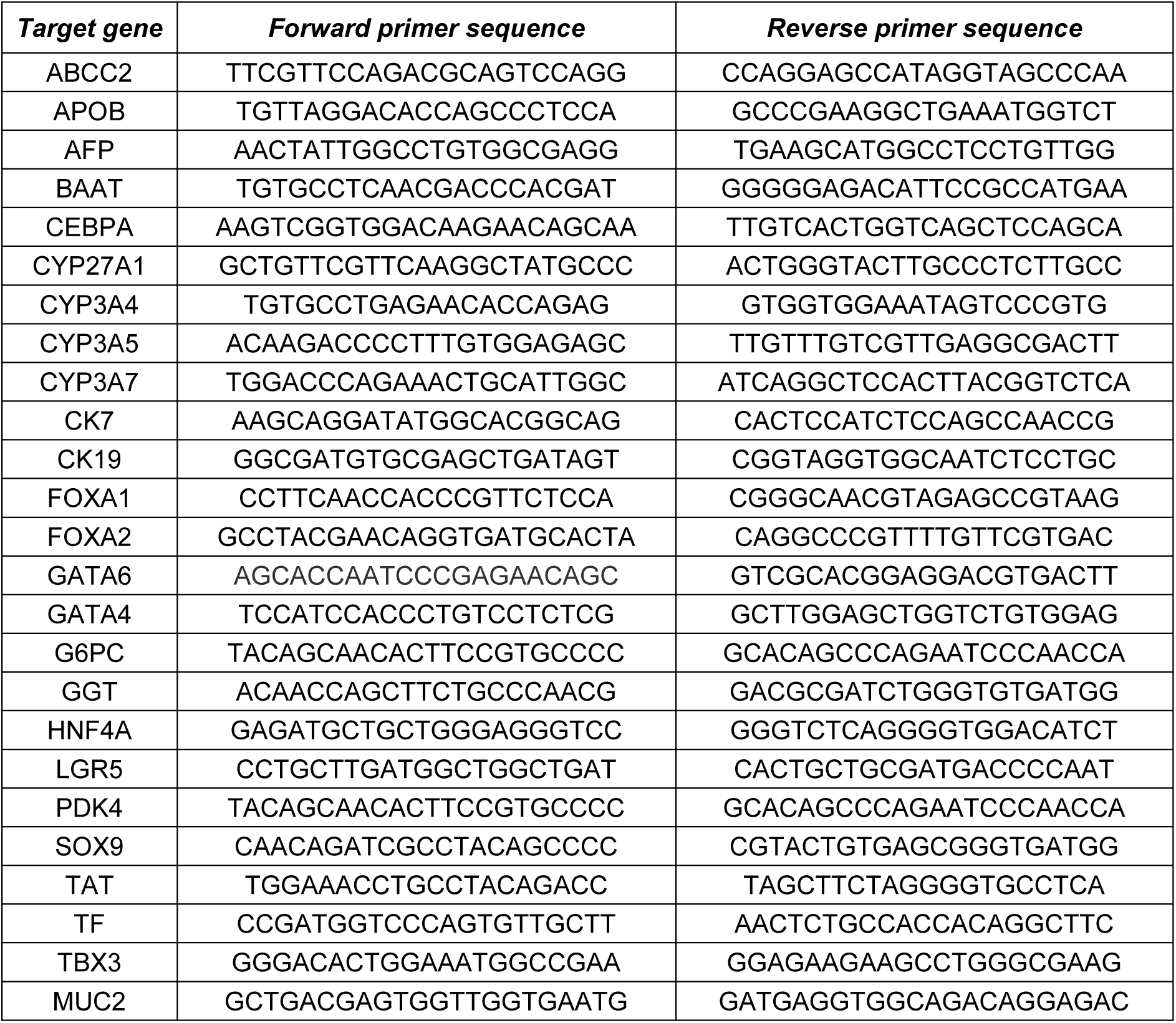
Sequences for forward and reverse primers used.

### 2.14 Enzyme-linked immunosorbent assays (ELISA)

Laminin concentrations in supernatants were measured using a laminin ELISA kit (Insight Biotechnology, #Boster Pikokine^TM^ EK0434). Acellular FLGs and fibrin gels were synthesised, and sterile DMEM/F12 media was added 30 min after gelation. Media was collected the next day (Day 1) and every other day for two weeks. Supernatants were stored at-20 °C until they were analysed alongside stock concentrations of laminin for comparison using the Laminin ELISA kit per the manufacturer’s instructions. Sample absorbance readings were then taken using a FLUOstar® Omega plate reader (BMG Labtech Ltd, UK) at 450 nm. The following equation estimated the residual amount of laminin in FLGs:

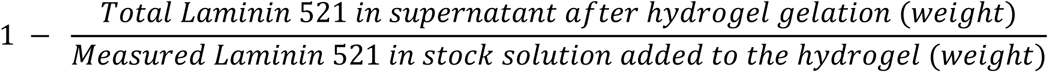

Albumin concentrations in supernatants were measured using a human albumin ELISA quantification kit (Bethyl Lab, #E80-129). Sample absorbance was measured with a Synergy HTX microplate reader (BioTek, USA). Supernatants from primary human hepatocytes (PHH) were collected 24 hr after the completion of cell recovery. Supernatants from iHeps were collected 2 days after (Day 22) the end of the differentiation protocol (Day 20); media were refreshed every other day as per the differentiation protocol.

Apolipoprotein B (APOB) secretion in the supernatant was quantified using the human APOB ELISA quantification kit (Mabtech, # 3715-1H-6) according to product literature. Sample absorbance was measured with an Infinite M1000 PRO microplate reader (Tecan, Switzerland). Supernatants from iHeps were collected 2 days after (Day 22) the end of the differentiation protocol (Day 20); media were refreshed every other day as per the differentiation protocol.

### 2.15 Functional Assays

#### 2.15.1 Cytochrome 3A4 (CYP3A4) assay

CYP3A4 activity was measured using the P450-Glo^TM^ CYP3A4 Assay kit (Promega, #V9001) as per the manufacturer’s protocol. Fluorescence in samples was measured using the Synergy HTX microplate reader. PHH after 24 hr of cell recovery, and Day 22 iHeps were used.

#### 2.15.2 Low-density Lipoprotein (LDL) uptake cell-based assay

LDL uptake was demonstrated using a cell-based kit (Cayman Chem, #10011125) following the manufacturer’s protocol. iHeps were incubated for 12 hr, then the HCM media was replaced with PBS. Images were then acquired immediately. Day 22 iHeps and hPSCs (negative control) were used.

#### 2.15.3 5-(and-6)-Carboxy-2’,7’-Dichlorofluorescein Diacetate (6-CDF)

6-CDF was used to demonstrate hepatocyte function following a protocol previously described (35)3D iHeps (Day 22) were incubated for 2 hrs in HCM media containing 1 μmol/L 6-CDF (Thermo Fisher, #C-369). The samples were then washed with PBS for 15 min before staining with Hoescht in HCM.

#### 2.15.4 Gamma-glutamyl Transfer (GGT) assay

GGT activity was determined using a fluorometric GGT assay kit (Abcam, #ab241030) according to the product literature. Fluorescence in samples was measured with the Infinite M1000 PRO microplate reader. PHH after 24 hr of cell recovery, and Day 22 iHeps were used. PHH after 24 hr of cell recovery, and Day 22 iHeps were used.

### 2.16 Animal experimentation

#### 2.16.1 Biocompatibility studies in mice

Biocompatibility studies were performed in immunocompetent general-purpose C57BL/6 (“Black 6”) mice obtained from InVivos (www.invivos.com.sg). Healthy, male mice between two and three months old (∼ 30 g to 35 g) were used. 100% croton oil (Merck, #C6719-10G) was emulsified in corn oil to a final concentration of 1.0% *v/v, referred to herein as “croton oil”.* Croton oil was used as a positive control because it is a well-cited skin allergen that is commonly used to study inflammation in murine models (36). 200 μl of croton oil (n=9), Alphagel (n=9), fibrin-only gel (n=9), and 0.9% saline (n=9) was injected subcutaneously using 18G hypodermic needles, and 1 ml low dead volume syringes into the flanks or scruffed backs of Black 6 mice with aseptic techniques. These volumes were within the internationally recommended limits for subcutaneous injections in mice (American Association for Laboratory Animal Science). Three mice from each group were culled at 1, 2, and 6 weeks post-injection. Mice were culled by cervical dislocation under general anaesthesia with isoflurane. Injection sites were excised and fixed in 10% formalin for 24 hr then transferred into 70% ethanol. Tissues were then embedded in liquid paraffin, sectioned by microtome, and placed on glass slides. Slides were stained with hematoxylin and eosin (H&E) to evaluate signs of inflammation, then imaged with the TissueFAXS i-plus system (TissueGnostics, Vienna, Austria).

#### 2.16.2 Hepatocyte transplantation in mice

NSG^TM^ immunocompromised mice (Jackson Laboratory, Invivos Singapore) were used to minimise the immunogenicity of xenotransplantation. For stem cell-derived hepatocytes in Hepatogel (H-iHep) transplantations, 3 x 10^6^ freshly harvested Day 22 H-iHeps were harvested and re-suspended in 75 μl of Hepatogel, then injected under direct vision with a 1 ml low dead volume syringe and 18G hypodermic needle into the inferior margins of the right liver lobe of the mice (n=9). As a negative control, stem cell-derived hepatocytes in Matrigel (i-Heps) were used. 3 x 10^6^ freshly harvested Day 22 iHeps were harvested, re-suspended in 75 μl of 0.9% saline, then injected under direct vision with a 1 ml low dead volume syringe and 18G hypodermic needle into the inferior margins of the right liver lobe of the mice (n=9). All mice received prophylactic antibiotics and analgesics subcutaneously for 3 days after the operation. Three mice from each group were culled on Day 3, Day 7, and 2 months post-hepatocyte transplantation.

Male and female NSG^TM^ mice were used. All mice were between four and five months old and weighed ∼ 40 g to 50 g. Mice were anaesthetised with isoflurane and had their abdomen shaved pre-operatively. Surgical laparotomies were performed aseptically to reveal the lower liver margin for injections.

All mice were fed a rodent maintenance diet (#2918 Teklad Irradiated Global, 18% Protein rodent diet, Envigo, Hackensack, NJ) and kept on BioCOB bedding. Approval for all animal work was granted by the National University of Singapore Institutional Animal Care and Use Committee (IACUC), protocol number R20-0822.

### 2.17 Statistical analyses

Prism 9.3.0 (GraphPad, USA) was used for data analysis and bar graph illustrations. Data were tested for normality using the Shapiro-Wilk test. Parametric data and their bar charts were presented as mean and standard deviation unless otherwise stated. For comparison between two groups of parametric values, the Student’s t-test was used to compare means. For comparing three or more groups of parametric data, one-way analysis of variance (ANOVA) with Tukey correction was used. For nonparametric data, the Chi-squared test or the Mann-Whitney U test was used to test for association; the Kruskal-Wallis test was used to compare three or more groups. All tests were two-sided, and the alpha was set at 5%. For the animal experimentation, the study had an 80% power to detect an effect size of 3.0. The TBtools-II software (37) was used to generate the Heat Map; data were normalised to hPSCs.

### 2.18 Ethical Approval

All human cell lines used are classed as non-relevant material under the Human Tissue Act 2004 (UK). As such, ethical approval was not needed. The use of animals and commercially obtained PHH was granted by the National University of Singapore (protocol number R20-0822).

## 3. RESULTS AND DISCUSSION

### 3.1 Focused screening revealed Laminin 521 and Fibrinogen as components in the embryological and hepatic niche that contribute to hPSC pluripotency

In previous work (38), iPSCs maintained on vitronectin in a defined feeder-free system were exposed to a library of proteins identified in the adult human liver by the Human Matrisome Project (http://matrisomeproject.mit.edu; Massachusetts Institute of Technology, USA). Hepatic matrix proteins that enhance the *in vitro* differentiation of iPSCs to iHeps when exposed to hepatic differentiation media (38) were then longlisted. However, to determine whether any of these hepatic proteins could maintain stem cell pluripotency, as observed during embryogenesis and liver organogenesis, we cultured hPSCs with these proteins in the absence of hepatic differentiation media. Successful candidates could then be used to biofabricate a hydrogel primed for liver tissue engineering when hepatic differentiation conditions are introduced into hPSC colonies. To that end, we optimised an automated high-throughput imaging pipeline to identify hPSCs by nuclear area and nuclear markers of stem cell pluripotency (NANOG, SOX2, and OCT4). Cell morphology and the mean fluorescent intensities (MFI) of pluripotency markers were used to define typical hPSC characteristics for positive selection (Supplementary Figure 1).

Using machine learning, the pipeline successfully identified pluripotent hPSCs as single cells growing in 2D colonies, with a sensitivity of 89.5% and a positive likelihood ratio of 160.8; Fig.1(B). Single-cell analyses excluded culture artefacts and cell clumps, which reduce the accuracy of automated image analysis. The pipeline identified non-pluripotent cell populations with a specificity of 99.4% and a negative likelihood ratio of 0.1; (Fig.1(C). The final readout of the pipeline was the Stemness Index (SI), a ratio of pluripotent cells to all DAPI-stained cells. The SI reliably discriminated between undifferentiated hPSCs and differentiated hPSCs within the same cell population; Fig.1(D). When tested, this platform demonstrated excellent discriminatory performance (Area Under the Curve (AUC) of 0.977); Fig.1(E).

Using this platform, we screened 22 proteins across >1,000 conditions and identified select proteins in both the embryonic and foetal liver niches that affect hPSC pluripotency. An example of a short-term (1-week) focused screening panel is illustrated in Supplementary Figure 2. In summary, the best candidate was chosen for its ability to maintain the highest SI in hPSCs after prolonged culture; Laminin 521 had the highest SI (between 0.95 and 1.0) after 3-4 months in culture. Thus, we sought to combine Laminin 521 with a polymeric backbone to form a 3D scaffold, since a pure or physiologically relevant Laminin 521 hydrogel is not currently available. We found that human fibrinogen was the most suitable polymeric protein to pair with Laminin 521 because of its mechanical tunability. At low concentrations, it did not cause rapid spontaneous differentiation of hPSCs (SI: 0.77-0.84). Notably, other polymeric backbones, such as albumin and collagen, either resulted in poor hPSC attachment or triggered significant spontaneous differentiation of hPSCs (SI < 0.5); examples are shown in Supplementary Figure 3.

### 3.2 Fibrinogen and Laminin concentrations inversely affected hPSC growth in 3D

To determine the optimum conditions for culturing hPSCs in 3D, we evaluated different compositions of Laminin 521-enriched fibrin hydrogels (FLGs) versus fibrin hydrogels (FGs) as controls. We found that both categories of hydrogels displayed distinctive characteristics. In 2D cell culture, cell attachment and expansion were significantly better in FLGs (70-100% confluence) versus FGs (10%-20% confluence) after 4-5 days in culture, Fig.2(A). This was reflected in the average fold increase in cell populations at 7 days (38.5 ± 5.6 vs 1.0 ± 0.2, p < 0.0001), respectively. In 3D cell culture, higher fibrinogen and lower laminin concentrations resulted in poor colony expansion and “dark”, unhealthy hPSC colonies (Fig. 2B), widely recognised features of spontaneous differentiation in hPSC culture (39–41). This was corroborated across several conditions by a significant decrease in SOX2, NANOG, OCT4 MFIs, and SIs compared to hPSCs grown on 2D Matrigel as standard practice (Supplementary Figure 4). Nonetheless, the optimal balance between fibrinogen, thrombin, and laminin concentrations was identified by combining SI in hPSCs and the stability of FLGs in cell culture; hydrogels with low fibrin concentrations (≤ 1mg/mL) disintegrated after a week in cell culture. This optimum ratio (50%:50% v/v Laminin 521 and 2.5mg/ml fibrin hydrogel) was termed “Alphagel”. Alphagel was found to be up to 3 times more cost-effective than Matrigel for 3D hPSC culture (Supplementary Figure 5). Unlike other compositions of FLGs, Alphagel supported the healthy expansion of hPSCs into 3D cell aggregates that closely resembled pluripotent spheroids (Fig. 2B). These aggregates contained fluid-filled cores; Fig.2(B) and Supplementary Figure 6. Cell viability assays (described below) indicated central necrosis was absent. In contrast, cell masses and liver organoids formed by suspension culture in media with low attachment surfaces, e.g. agarose coatings (42,43), tend to have cell-laden cores prone to central necrosis as these aggregates or organoids enlarge.

**Figure 2.**
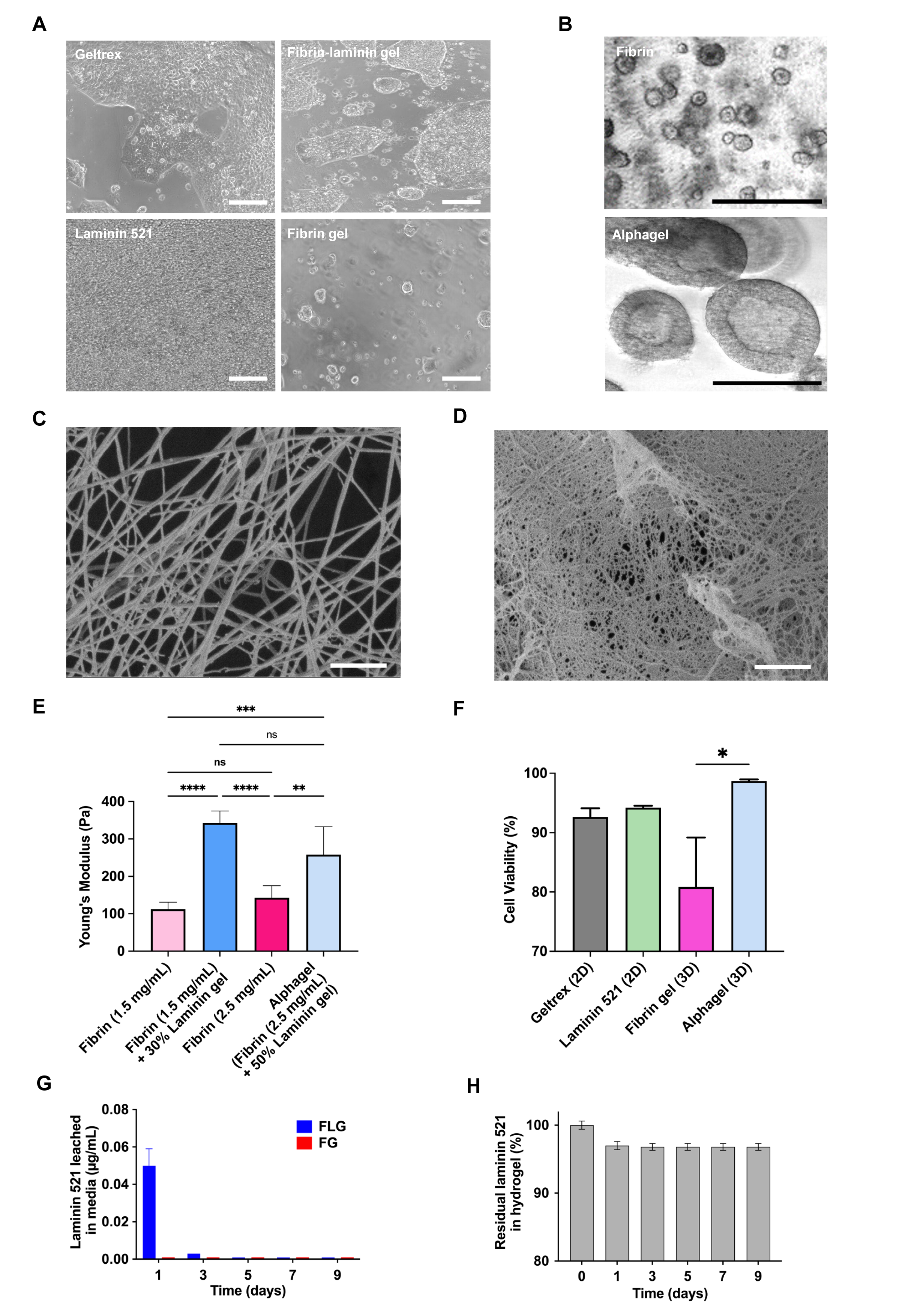
A) hPSC confluence in 2D growth factor reduced Matrigel (Geltrex), Laminin 521, fibrin-laminin hydrogel, and fibrin gel after 4 days. Scale bar = 100 μm. B) hPSCs cultured in 3D hydrogels. Top panel: Fibrin (5 mg/mL) gel. Bottom panel: Alphagel containing structures resembling pluripotent spheroids, scale bar = 200 μm. SEM of C) fibrin gel and D) Alphagel, scale bar = 1 μm, magnification 20K X, iProbe = 13 pA, 2.00 kV, WD = 4.4 mm for both images. E) Young’s moduli in hydrogels with varying fibrin and laminin concentrations. One-way ANOVA; ** = *p* < 0.01, *** = *p* < 0.001, and **** = *p* < 0.0001. F) Cell viability of hPSCs cultured in Alphagel, fibrin-only hydrogels, and 2D standard substrates. Mean ± S.E.M displayed. t-test; * = *p* < 0.05. G) ELISA of Laminin 521 in culture media used with acellular Alphagel and H) the calculated amount of fibrin-bound laminin over time.

To visualise the differences between fibrin-only hydrogels and Alphagel, scanning electron microscopy (SEM) was performed on acellular hydrogels. This demonstrated smaller pores and thinner fibrin fibres, as previously observed in laminin-fibrin hydrogels (32,44). Thin membranes in between inter-connecting fibrin fibres in Alphagel were observed, and these were absent in fibrin-only hydrogels; Fig.2(C) and Fig.2(D). As protein concentration alters the elastic properties of hydrogels, we next measured Young’s modulus of fibrin-only hydrogels, FLGs, and Alphagel. We found that hydrogel stiffness increased significantly with increasing laminin and fibrin concentrations (Fig. 2E). The Young’s modulus in FLGs containing 1.5 mg/ml fibrin was higher compared to 1.5 mg/ml fibrin-only hydrogels (fold increase 3.6 ± 0.6, p < 0.0001) and 2.5 mg/ml fibrin-only hydrogels (fold increase 2.6 ± 0.2, p < 0.0001). Despite the increase in stiffness, cell viability was notably higher in Alphagel compared to fibrin-only hydrogels (98.7 ± 0.3% vs 80.8% ± 8.3%, respectively, p <0.05); Fig.2(F). There were no significant differences in cell viability across Alphagel and the 2D substrates.

Next, to determine if the FLG hydrogels were stable, we measured the amount of Laminin 521 leached from acellular Alphagel into the culture media over time. This showed that the greatest loss of Laminin 521 from the hydrogel occurred during the first 24 hr of conjugation (approximately 3% leached into media). Thereafter, the leaching of Laminin 521 into the culture media was negligible, and approximately 97% of the laminin remained within the hydrogel after 9 days; Fig.2(G) and Fig.2(H). Then, to optimise 3D culture conditions, we tested the performance of hPSC-laden Alphagel paired with different stem cell maintenance media over 2 months. We found the best stem cell culture media that maintained markers of hPSC pluripotency with Alphagel were the TeSR-only family, i.e., TeSR^TM^ 2, mTeSR^TM^ 1, and mTeSR^TM^ Plus, yielding SIs for hPSCs of 0.95-1.0 (Supplementary Figure 4). Despite prolonged culture, pluripotency markers (OCT4, NANOG, and SOX2) were still strongly expressed in the 3D hPSCs bodies. For validation, these aggregates were dissociated into smaller cell clusters and cultured as a single-cell layer in 2D. Importantly, these hPSCs continued to proliferate in 2D and continued to express all 3 pluripotent nuclear markers (Supplementary Figure 6).

In summary, Alphagel is stable hydrogel that supports the formation and expansion of pluripotent spheroids in prolonged 3D hPSC culture, with distinct biological and physical properties compared to fibrin-only hydrogels. Thus, Alphagel provides a physiological environment to hPSCs because Laminin 521 and 511 are present in the inner cell mass of the blastocyst and its basement membranes during embryological development. (45,46). Its expression remains constant throughout inner cell mass expansion and during differentiation, whereas Laminin 511 expression decreases during differentiation (46). Importantly, through RNA sequencing (RNAseq) of human adult and foetal liver tissue, we found that Laminin 521 is differentially expressed in the maturing foetal liver (38), along with a small group of ECM proteins that also support stem cell pluripotency to some extent. Notably, Fibrinogen is also produced by trophoblasts during early development (47,48) and has the potential to be used as an injectable biomaterial, as it is abundantly present in human serum. Cell and protein adhesion is supported by arginine-glycine-aspartate (RGD) binding motifs at positions 96 and 572-574 on the α chains, heparin-binding domains (HBDs) also on the α chains, and a carboxy-terminal peptide (HHLGGAKQAGDV) at position 400-11 on the γ chains (49,50). Likewise, multiple binding sites on the α chains (RGD and HBD) and β chains (mainly RGD) in laminins facilitate cell and protein adhesion (51,52). These help explain the difference seen between Alphagel and fibrin-only gels.

Notably, there is conflicting data on the effects of fibrin hydrogels on stem cell culture. It was previously reported that fibrin gels (30 mg/ml) could be used for iPSC culture (53). Fibrin concentration is directly proportional to the stiffness of fibrin matrices (54), and the 10 mg/ml fibrin-only gels we tested had a Young’s modulus of ∼ 2 kPa. In our experiments, high concentrations of fibrin (≥ 4 mg/ml) promoted spontaneous differentiation across 3 hPSC cell lines (2 ESC cell lines and 1 iPSC cell line), and this was augmented by the type of stem cell maintenance media used (Supplementary Figure 4). This was evidenced by the low differentiation efficiency and suboptimal end-target cell phenotypes in fibrin-only hydrogels (reported below), and by independent studies showing that compliant substrates maintain ESC self-renewal by downregulating cell-matrix traction forces, whilst stiffer substrates promote spontaneous differentiation by downregulating OCT3/4 expression (55,56). In contrast, our FLGs were finetuned to a Young’s modulus of ∼ 250-400 Pa to mimic the stiffness of embryological liver. This range was determined by atomic force microscopy of foetal liver tissue in other studies (57,58), which also showed that compliant matrices facilitated liver tissue expansion better than stiffer substrates. Interestingly, it was previously demonstrated that compliant fibrin-only gels (1 mg/ml, ∼ 100 Pa) promoted the selection and growth of tumour cells that express stem cell markers CD133, Nestin, BMI-1, and C-kit, but not stem cell markers OCT3 and NANOG (56). In our study, adding Laminin 521 to low-concentration fibrin matrices preserved high OCT4 and NANOG expression in hPSCs over prolonged culture (2-3 months). Similarly, fibrin-only hydrogels resulted in the absence of OCT3 and NANOG expression.

### 3.3 Alphagel supported the tri-lineage differentiation of hPSC bodies into cardiac, neural, and hepatic tissue in 3D

To rigorously demonstrate that hPSCs cultured in Alphagel remained pluripotent, we cultured hPSCs bodies in Alphagel, then performed directed 3D differentiation of hPSCs into all three germ cell lines: mesoderm (cardiac), ectoderm (neurons), and endoderm (hepatic).

During cardiac specification, hPSC bodies in Alphagel successfully proliferated and formed 3D contractile cardiac tissue after initiating cardiac differentiation; Fig.3(A) and Supplementary Movie 1. Immunofluorescent staining (IF) of cells for the cardiac marker Troponin T (TnT) and the general myoblast marker α-SMA (59) was strongly positive; Fig.3(B). Cardiomyocytes began beating at 30-50 beats/min between Days 5 and 8 of the cardiac differentiation process; early contractions were weak and asynchronous (Supplementary Movie 1). However, the contractility rate increased to 60 ─ 80 beats/min, and contractions were noticeably stronger and synchronous by Day 12 ─ 14 (Supplementary Movie 2). There was no significant difference in differentiation efficiency (proportion of TnT positive cells) between Alphagel and Matrigel (75.4 ± 18.5% vs 78.2 ± 15.4%, respectively, p = 0.14). In fibrin-only hydrogels, the differentiation efficiency was significantly lower versus Alphagel (34.9 ± 21.3%; p < 0.001); Supplementary Figure 7(A).

**Figure 3:**
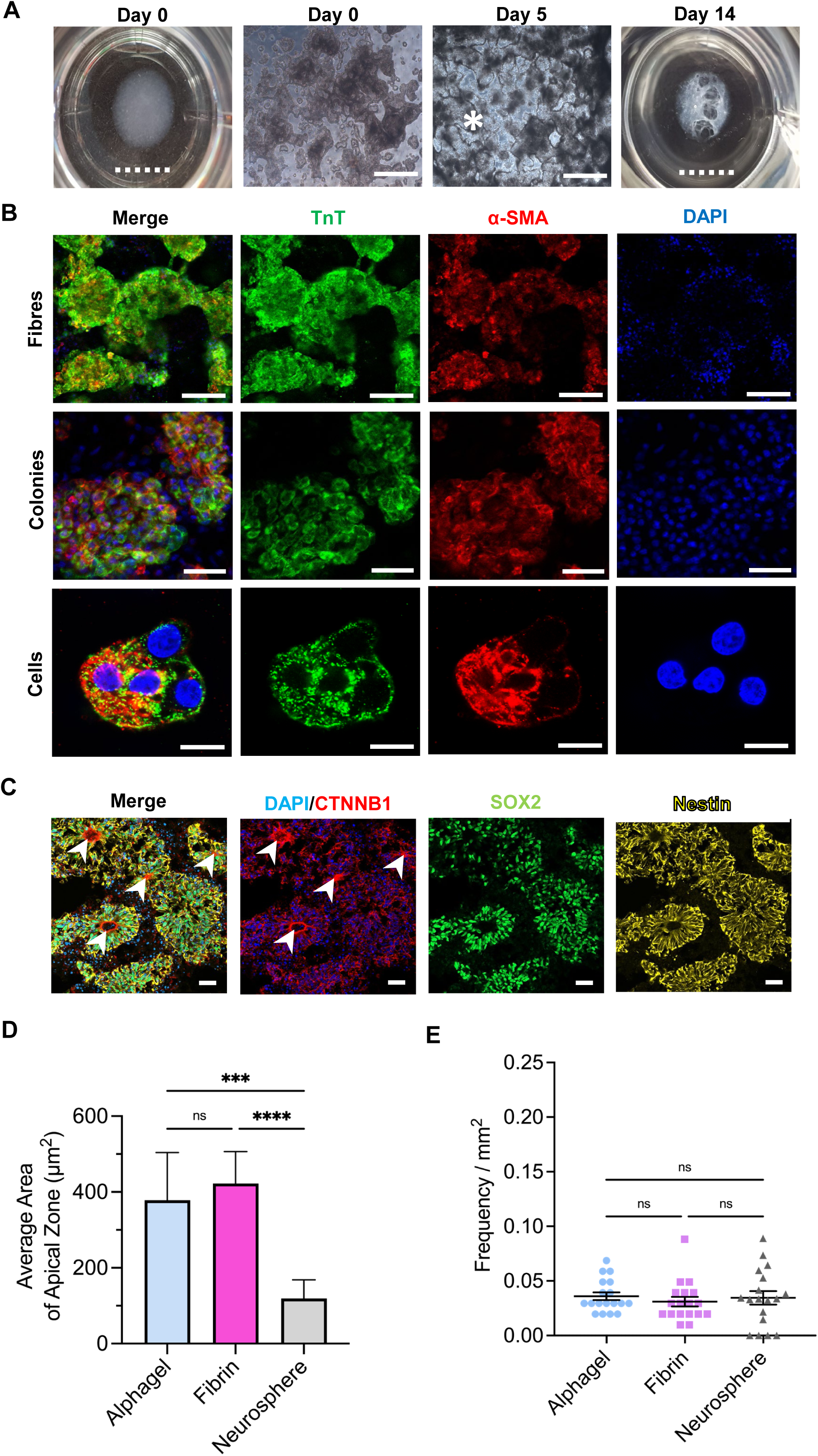
A) Macroscopic and microscopic appearances of cells in Alphagel. Contractile cardiomyocyte colonies (*) are visible by Day 6. By Day 14 contractile tissue resembling trabeculae carnae is observed. Scale bar: full line = 100 μm, dotted line = 1 cm. B) Troponin T (TnT), alpha-smooth muscle actin (α-SMA), and DAPI in cardiomyocytes cultured with Alphagel. Scale bar = 100 μm (fibres), 50 μm (colonies), and 20 μm (cells). C) Alphagel-derived neuroepithelia expressing β catenin (CTNNB1), SOX2 and Nestin. Arrows indicate apical zones demarcated by β catenin. Scale bar = 100 μm. D) Average lumen area of neuroepithelia in Alphagel, fibrin gel, and neurospheres (Matrigel). E) Frequency of polarized puncta per mm^2^. The Kruskal-Wallis test was used for D) and E). *** = *p* < 0.001, and **** = *p* < 0.0001.

For neural specification, hPSCs were subjected to a commonly applied protocol utilising dual-SMAD inhibition (60,61). For comparison, both a fibrin-only control and an aggregate-based (neurosphere) method were included. Neurospheres are derived from hPSCs cultured with Matrigel and then differentiated in suspension culture. At Day 16, all conditions showed successful acquisition of a neural fate as demonstrated by co-expression of the neuroepithelial progenitor markers SOX2, nestin, and β-catenin (62); Fig.3(C). Quantifications at this time point showed comparable cell composition of SOX2 immunoreactive (IR) progenitor cells in Alphagel (79.5 ± 5.0%) and neurospheres (82.4 ± 9.3%) conditions. In the fibrin-only condition, there were slightly fewer SOX2 IR cells (74.1 ± 12.6%); however, this difference was not statistically significant; Supplementary Figure 7(B). To determine whether the structural properties of Alphagel could improve the organisation and morphology of differentiated neuroepithelia, we counted and measured the number and size of lumens demarcated by nestin/β-catenin co-labelled apical cells and compared them with those observed in fibrin-based hydrogels and neurospheres. We found that the frequency of polarised areas (approximately 0.03/mm^2^) was consistent across all conditions, albeit less variable under hydrogel conditions; Fig.3(D). However, the average lumen area was significantly increased in hydrogel conditions with Alphagel (403.29 µm^2^; p = 0.0007) versus neurospheres (105.46 µm^2^); Fig.3(D). Large lumen areas are observed when neural progenitors coalesce and mature (63,64), forming the structural architecture of the neural progenitor niche. The proper organisation of polarised tissue is crucial for neurodevelopment, and aberrant formation is implicated in neural tube defects. Nonetheless, no differences were observed in the frequency of polarised puncta between Alphagel and neurospheres (Fig. 3E). Cumulatively, these results show that Alphagel supports efficient ectodermal differentiation of hPSCs into neuroepithelia.

For hepatic specification, successful differentiation of hPSCs into hepatocytes (iHeps) was demonstrated through IF staining, showing previously established gene expression signatures (35), and functional assays. Positive IF staining for hepatocyte markers Albumin, HNF1A, HNF4A, CYP2A6, CD147, and E-cadherin was demonstrated in 3D-differentiated cells, Fig.4(A). Similarly, key hepatocyte genetic markers HNF4A, CEBPA, and CYP3A4 were shown to be expressed in iHeps derived in Alphagel at levels similar to those in iHeps derived in Matrigel Fig. 4(B). However, CYP3A7 and AFP expression were significantly higher in Alphagel and Matrigel than in adult primary human hepatocytes (PHH); p < 0.0001 for both, Fig. 4(B). Notably, TBX3 expression was also high in Alphagel and Matrigel, but there was no statistically significant difference compared to PHH. For further characterisation, 18 other hepatocyte-related genes were analysed (Supplementary Figure 8). 10 of 18 genes showed differences between PHH and Alphagel-derived hepatocytes. Specifically in the latter, BAAT, ABCC2, and TAT were down-regulated, whilst CYP3A5, GATA4, GATA6, FOXA1, SOX9, LGR5, and CK19 were up-regulated compared to PHH (p values in Supplementary Figure 8). BAAT and TAT code for liver enzymes, and ABCC2 is a cell-membrane transporter. CYP3A5 codes for a hepatic enzyme that increases as the foetal liver matures. GATA4, GATA6, and FOXA1 are important transcription factors involved in hepatic differentiation. LG5R and SOX9 are markers for hepatic progenitors (hepatoblasts) that give rise to the two main cell types within the liver (hepatocytes and cholangiocytes). CK19 is a biliary marker.

**Figure 4:**
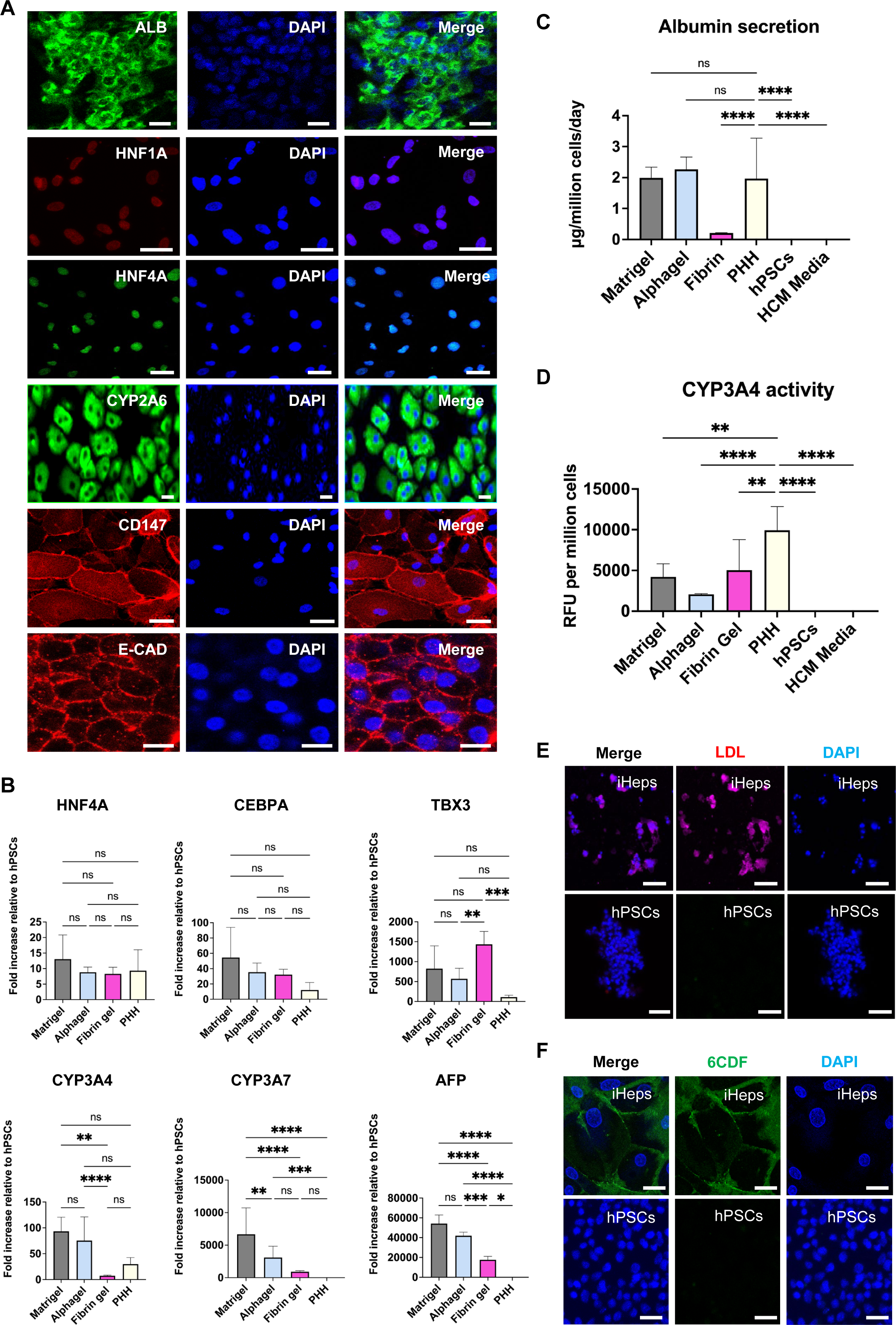
A) Key hepatocyte markers in Alphagel-derived iHeps. ALB = albumin, HNF = Hepatocyte Nuclear Factor, CYP2A6 = Cytochrome 2A6, CD147 = Cluster of Differentiation protein 147, and E-CAD = E-cadherin. Scale bar = 25 μm. B) Key hepatocyte markers by gene expression (qPCR). C) Albumin secretion (ELISA) and D) CYP3A4 activity (P450-Glo^TM^) in PHH vs iHeps cultured in various gels (Day 22). HCM = Hepatocyte Culture Media (Lonza). E) LDL uptake (red) in Alphagel-derived iHeps versus hPSCs. Scale bar = 100 μm. F) CDFDA secretion (green) in Alphagel-derived iHeps vs hPSCs. Top panel scale bar = 20 μm, bottom panel scale bar = 50 μm. One-way ANOVA was used; * = *p* < 0.05, ** = *p* < 0.01, *** = *p* < 0.001, and **** = *p* < 0.0001.

To assess the functionality of iHeps derived in Alphagel, albumin secretion, cytochrome 3A4 activity, LDL uptake, as well as CDFDA uptake and excretion were investigated at the end of the hepatic differentiation process. Albumin, a key marker of hepatocyte synthetic function (65), was found to be secreted in similar amounts by PHH and iHeps cultured in Alphagel and Matrigel; Fig.4(C). However, CYP3A4 activity was lower in Alphagel and Matrigel than in PHH, as reported by other groups (35,66); Fig.4(D). Importantly, LDL uptake and CDFDA secretion, both features of hepatocyte metabolic function, were demonstrated in Alphagel; Fig.4(E) and Fig.4(F). These were also present in primary hepatocytes and hepatocytes derived in Matrigel. There was no statistically significant difference in differentiation efficiency between Alphagel and Matrigel (defined herein by the proportion of albumin-positive cells). This was approximately 70.6% and 69.9%, respectively (p=0.73). In fibrin-only hydrogels, the differentiation efficiency was significantly lower versus Alphagel (28.8%, p<0.0001); Supplementary Figure 7(C).

In summary, the results above show that Alphagel maintains hPSCs pluripotency and supports their trilineage differentiation in 3D. Cardiac, neural, and hepatic tissue were derived in 3D and positively characterised using established and chemically defined differentiation protocols. We did not perform a teratoma formation assay because expert consensus favours tri-lineage differentiation as a demonstration of pluripotency; teratoma formation assays yield inconsistent results (67). In our experiments, differentiation efficiency varied between hPSC cell lines. However, these were similar to the reported literature (68–70). Nonetheless, the organisation and polarisation of tissues are important to tissue function. In particular, this has been a defining feature previously optimised in neural organoid protocols (71). While these methods typically rely on embedding early aggregates in Matrigel, we show that Alphagel has similar properties, allowing the spontaneous development of organised neural tissue with enlarged apical lumens. Besides consistency, a significant advantage of the Alphagel system is that it allows simultaneous differentiation and structural organisation without the need for further physical manipulations or tedious time-consuming steps, and thus could be helpful to explore as a substrate for brain organoid generation. Similarly, we show that the platform enables the generation of 3D cardiac tissue without requiring substrate changes mid-differentiation, thereby holding promise for the development of injectable regenerative therapies for cardiac disease or for ex vivo cardiac tissue engineering.

### 3.4 Alphagel demonstrated good biocompatibility and biodegradability *in vivo*

To determine whether Alphagel could be used as a biomaterial for clinical therapy, we next assessed its biocompatibility *in vivo*. To that end, we injected immune-competent Black 6 (C57BL/6) mice subcutaneously with Alphagel (test group), 0.9% saline (negative control), and 1.0% v/v croton oil (inflammatory agent, positive control) to assess for an immune response; n=3 for each group and time point. Mice were then observed and culled over 6 weeks. Histological slides were evaluated by a veterinary histopathologist who was blinded to the groups. Each slide was scored objectively using an international scoring tool (Supplementary Figure 9) and graded using standardised nomenclature (72–74). The international scoring tool provided a quantitative measure of key aspects in inflammation, e.g. cell infiltrates, fibrosis, etc.

Cumulatively, the results showed that mice injected with Alphagel had mild inflammation at the injection site between 1 and 2 weeks; Fig. 5(A). However, this resolved completely between 4 and 6 weeks; Fig.5(B). Mice injected with 0.9% saline also showed minimal inflammation at the injection site after 1 week, which resolved completely by 2 weeks. Mice injected with 1% croton oil developed ongoing severe inflammation, which progressed to wound dehiscence by 6 weeks; these mice were culled humanely when wound inflammation became visibly severe. The histological scores were consistent with the qualitative assessment of the histological slides (Fig. 5C).

**Figure 5:**
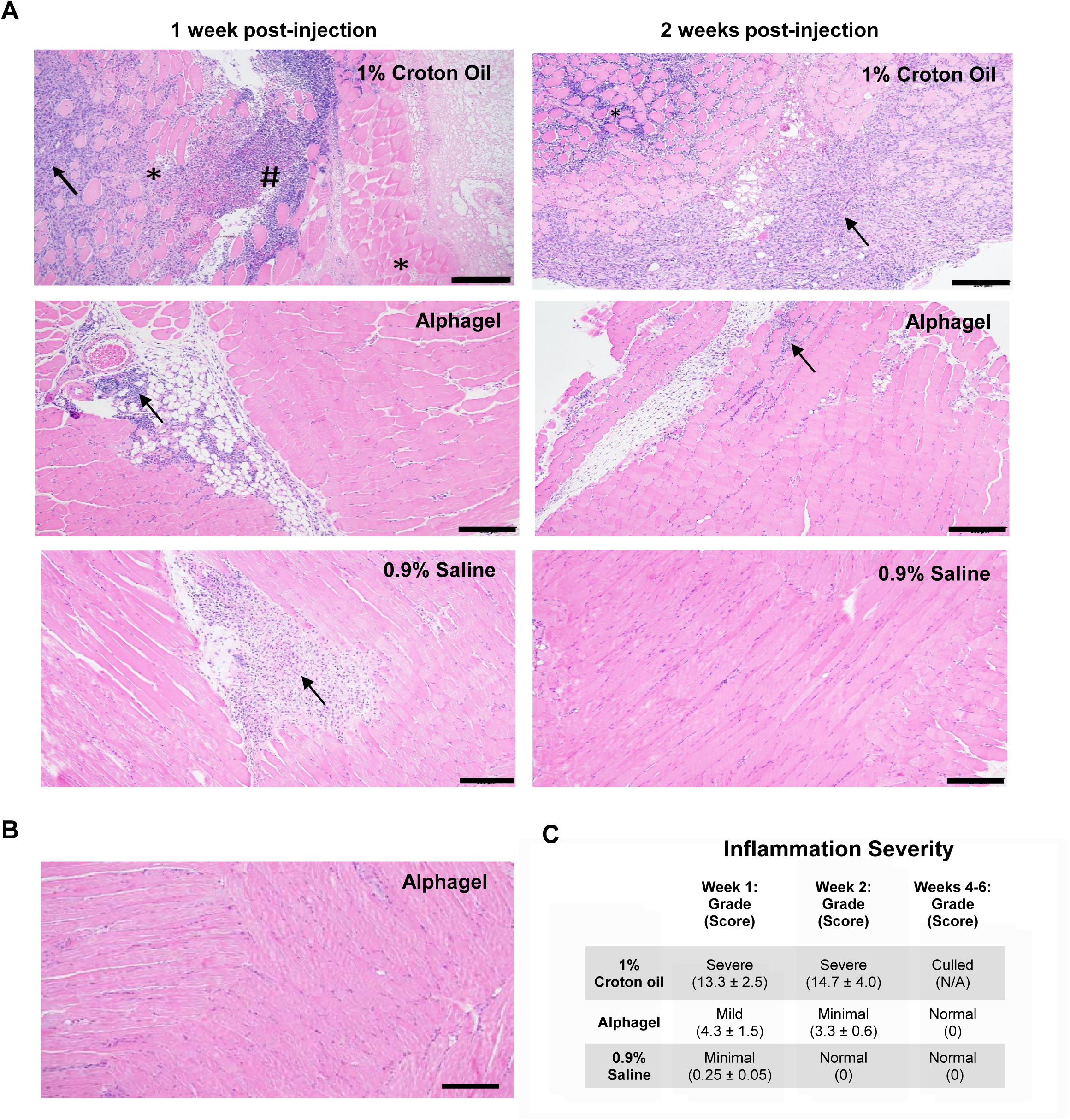
A) H&E stains of sites injected with 1% Croton Oil, Alphagel, and 0.9% Saline at 1 and 2 weeks. Black arrow denotes areas of granulation and inflammation, * denotes necrotic and degenerating muscle fibres, and # denotes supportive changes around affected muscle fibres. Scale bar = 200 μm. B) H&E stain of Alphagel injection site after 6 weeks showing complete resolution of previously mild inflammation. Scale bar = 200 μm. C) Histological grade and scores of injection sites over 6 weeks. One-way ANOVA; * = *p* < 0.05, ** = *p* < 0.01, *** = *p* < 0.001, and **** = *p* < 0.0001.

Neoplasia was not observed at the injection sites in all groups, and Alphagel was not visible at the injection sites between 4 and 6 weeks. It is noteworthy that the degradation of Alphagel (and all FLGs) is dependent on the presence of fibrinolytic enzymes such as plasmin. In vitro, acellular Alphagel remained stable for over two months. Cellularised Alphagel hydrogels were resistant to chemical dissociation, unlike cellularised 2D substrates, which typically dissolve readily (Supplementary Data). Notably, cells embedded within the hydrogels and in vitro conditions will influence the production of fibrinolytic enzymes and the rate of degradation, which can range from 2 to 6 weeks. In summary, data from immunocompetent Black 6 mice demonstrate that fibrin-laminin hydrogels are biocompatible and biodegradable *in vivo*. The mild local inflammation in immunocompetent Black 6 mice was likely a combination of iatrogenic injury (given similar reactions to saline injections) and the implantation of xenogeneic biomaterials (human ECM proteins), suggesting it would be well tolerated in human studies.

### 3.5 Alphagel enriched with liver-specific ECM (Hepatogel) enhanced the phenotype of human iHeps

To test our hypothesis that hydrogels enriched with organ-relevant ECM can improve stem cell-derived phenotypes, we added Laminin 411 and Laminin 111 to Alphagel. These were specific ECMs identified as important in liver development (38). Various combinations of protein concentrations were tested using functional assays (Supplementary Figure 10), and the concentrations of Laminins 521, 411, and 111 that yielded the best results were termed “Hepatogel”. iHeps derived in Hepatogel, Alphagel and Matrigel were compared against PHH using the same panel of liver-specific genes and functional assays. Key changes in the gene expression profile are highlighted in Fig.6(A). A heat map illustrating the differential gene expression across the 24-gene liver panel is provided Fig.6(B); statistical comparisons are better illustrated in Fig 6(A) and Supplementary Figure 11. Briefly, Hepatogel increased the gene expression for the hepatic glycoprotein Transferrin (TF) versus Alphagel (1.5 ± 0.4 fold increase, p < 0.0001) and PHH (2.2 ± 0.6 fold increase, p < 0.001). The gene expression of the liver enzyme Gamma Glutamyl-Transferase (GGT) was also significantly increased across all three hydrogels compared to PHH, but it was highest in Hepatogel (3.3 ± 0.3-fold increase vs PHH, p < 0.05). Notably, gene expression of the liver enzyme Glucose-6 Phosphatase (G6PC) was significantly lower in Matrigel versus PHH (-6.7 ± 0.6 fold, p < 0.05); however, expression in Hepatogel was comparable to PHH. Bipotent-hepatoblast marker LGR5 was significantly upregulated in Alphagel (6.0 ± 0.6 fold increase; p<0.01) and Hepatogel (6.5 ± 0.5 fold increase; p < 0.01) versus PHH. Importantly, expression of immature foetal liver markers CYP3A7 and AFP were significantly lower in Hepatogel versus Matrigel (1.8 ± 0.4 fold decrease; p < 0.05 and 1.3 ± 0.1 fold decrease; p < 0.05, respectively).

**Figure 6:**
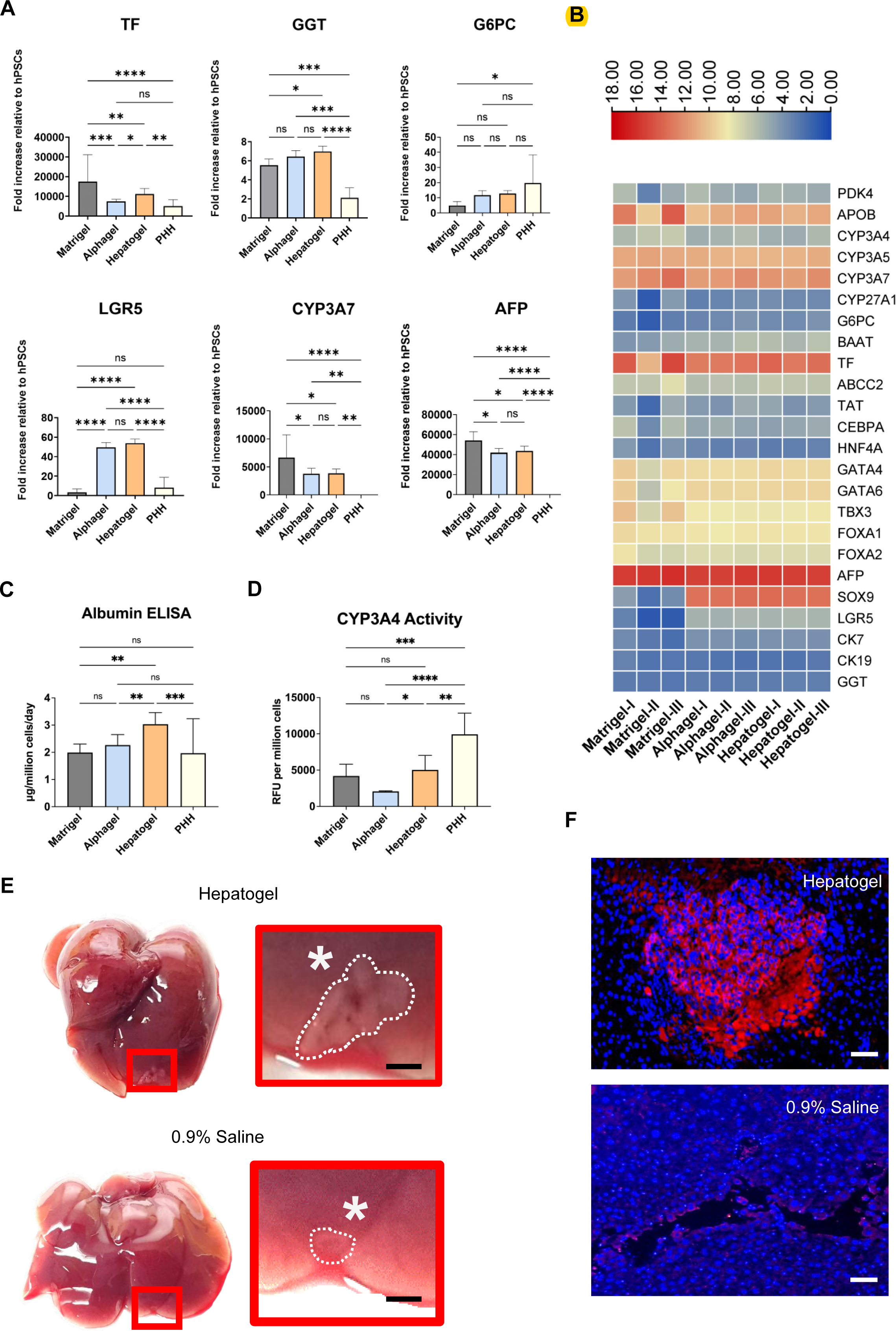
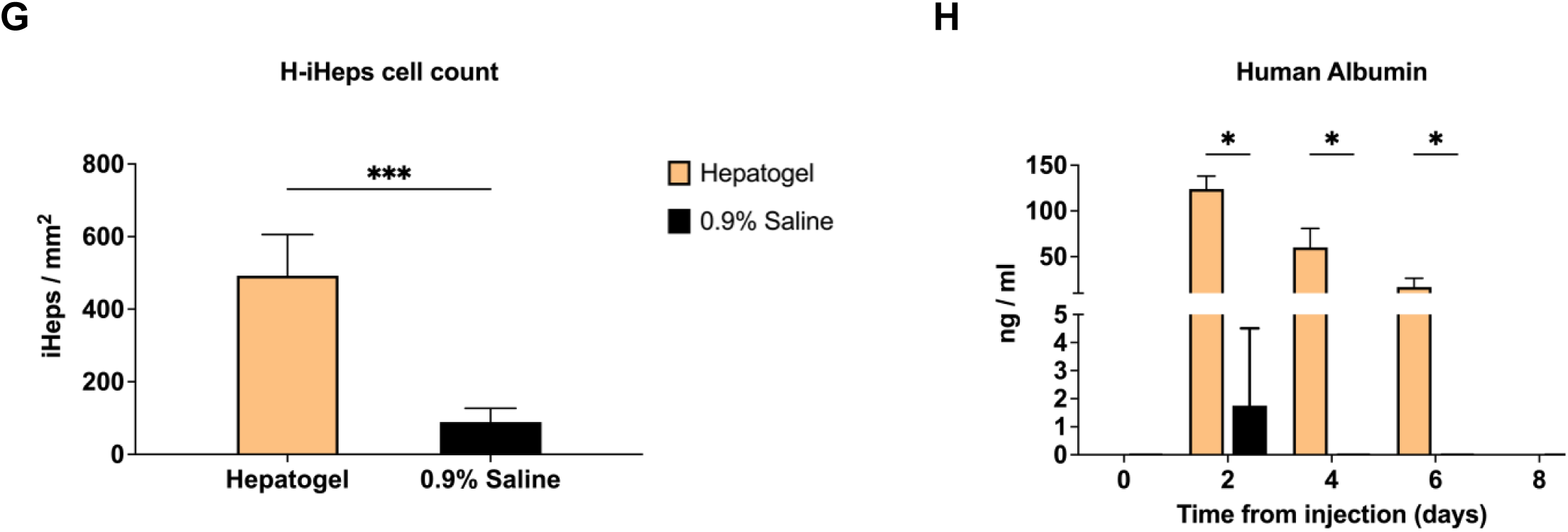
A) Differentially expressed genes in iHeps; Hepatologel, Alphagel, Matrigel, and adult primary human hepatocytes (PHH). B) A heat map summarising the differential gene expression of a hepatic 24-gene panel across replicates of Matrigel, Alphagel, and Hepatogel (normalised to hPSC). C) Albumin ELISA of culture media and D) Luciferin-based measure of CYP3A4 activity; 2 days after completion of iHep differentiation and 2 days after plating PHH. One-way ANOVA; * = *p* < 0.05, ** = *p* < 0.01, *** = *p* < 0.001, and **** = *p* < 0.0001. E) Mouse livers 3 days after intra-hepatic injection with H-iHeps in Hepatogel and 0.9% saline. Red box = area magnified. * = site of injection, dotted white lines demarcate engrafted cell mass, scale bar = 1 mm. F) Human albumin (stained red) in engrafted iHeps 3 days after intra-hepatic injection. Scale bar = 100 μm. G) H-iHeps identified by albumin staining on liver histology 3 days after intra-hepatic injection. H) ELISA of mouse serum for human albumin after injection with H-iHeps in Hepatogel and 0.9% saline over time. Day 0 = serum levels before injection. T-test; *** = *p* < 0.001, and **** = *p* < 0.0001.

The synthetic function of hepatocytes was assessed by albumin secretion. Hepatogel induced the highest albumin secretion compared to Alphagel, PHH, and Matrigel; Fig.6(C). The metabolic function of hepatocytes was assessed by CYP3A4 activity and APOB secretion. CYP3A4 activity was highest in PHH. iHeps derived in Hepatogel had higher CYP3A4 activity than Alphagel (2.4 ± 0.1 fold increase; p < 0.05) and Matrigel. However, in the latter, the increase was marginal and not statistically significant (1.2 ± 0.5; p = 0.47), Fig.6(D). APOB secretion was higher in Hepatogel than in PHH; however, there was no difference between Hepatogel and Alphagel (Supplementary Figure 12). In summary, the two liver-relevant ECMs added to Alphagel increased TF expression, CYP3A4 activity, and albumin secretion compared with Alphagel alone. Interestingly, up-titrating Laminin 411 concentrations in Hepatogel significantly increased APOB secretion. Conversely, this promoted the formation of branching 3D tubular structures lined by cells that expressed biliary markers CK7, CK19, and SOX9 (Supplementary Figure 13, Supplementary Movie 3, and Supplementary Movie 4), accompanied by a corresponding increase in Alkaline Phosphatase (ALP) in the culture media (Supplementary Figure 12).

The concentration of Laminin 411 in Hepatogel likely led to modest improvements in hepatocyte phenotype compared with Alphagel and Matrigel. This was evidenced by significantly higher albumin expression paired with lower foetal liver markers CYP3A7 and AFP. Integrin pathways offer a credible mechanism. The C-terminal of the Laminin 411 domain binds α3β1 integrin receptors, critical mediators of hepatocyte differentiation (75,76). α3β1 integrin receptors are highly expressed on immature hepatocytes, transformed hepatocytes, and biliary cells. Its binding triggers the activated protein kinase/extracellular signal-regulated kinase (MAPK/ERK) (77,78), focal adhesion kinase (FAK) (79,80), and Yes-associated protein (YAP) (81) in epithelial cells. MAPK/ERK signalling activates the pregnane X receptor (PXR) (82), which in turn induces CYP3A4 activity (83). Similarly, MAPK/ERK signalling via Runx1 can activate CCAAT/enhancer-binding protein alpha (CEBP/a) gene expression, which is responsible for albumin synthesis (84,85). The FAK/YAP pathway may have also resulted in the formation of biliary structures within the liver tissue (86). Though interesting, demonstrating the mechanistic sequelae of Laminin 411 binding to integrin receptors is beyond the scope of this study.

Nonetheless, it likely that a synergy between Laminin 521, Laminin 111, and Laminin 411 exists. All three laminin isoforms are differentially expressed (on RNAseq) at various stages of liver development (37,86,87) and appear to play distinct roles in liver development. Laminin 521 and Laminin 511 are abundant in the basement membranes of developing and regenerating livers. Notably, a combinatorial effect between Laminin 521 and Laminin 111 has been shown in hepatocytes, but the exact mechanism by which this occurs is unclear (87). On its own, Laminin 111 promotes the differentiation of hPSC into primitive endoderm (25–27,88), maintains hPSC-derived bipotent hepatoblasts in 3D culture (89,90), and controls apicobasal polarisation of liver epithelial cells in the developing liver, before being downregulated in healthy adult livers but upregulated in the presence of liver injury (91). Separately, Laminin 411 augments the effects of ECM proteins such as Laminin 111 and Laminin 521. The binding of Laminin 411 to α3β1 integrin receptors induces morphological changes in immature and transformed hepatocytes, enabling them to interact more effectively with spatial ECM components, thereby enhancing differentiation (75). Single-cell spatial transcriptomics of human livers has shown that genes coding for the Laminin α4 (LAMA4) and β1 (LAMB1) chains are differentially upregulated in hepatic stellate cells, liver sinusoidal endothelial cells, zone 1 and 3 hepatocytes, and some cholangiocytes (92). This distinguishes Laminin 411 from Laminin 521 and Laminin 111, especially in cells in the periportal area (zone 1), where LAMA4 and LAMB1 are highly expressed, suggesting that Laminin 411 plays an important role in hepatic zonation. Notably, the LAMA4 chain critically regulates endothelial specification and the maturation of capillaries, veins, and arteries. It influences cell morphology and migration, thereby contributing to the formation of 3D structures, such as early bile ducts. In contrast, the LAMA5 chain is present in the basement membrane of veins and capillaries, but not in arteries (93,94). These complex relationships between the laminin isomers help explain the effects of Hepatogel and demonstrate the importance of multi-protein substrates for in vitro liver organogenesis.

### 3.6 Hepatogel improved the engraftment of transplanted hepatocytes compared to standard therapy

Since iHeps cultured in Hepatogel (H-iHeps) had a better synthetic (protein synthesis) and metabolic profile than Alphagel, we next investigated if, by introducing a proliferative/regenerative liver niche at the site of cell transplantation, the retention of transplanted hepatocytes could be improved. At present, human hepatocytes are commonly delivered in solution via injections, either directly into the liver or into various vascular sites that feed into the liver, for cell therapy (95). However, in these approaches, cell retention and long-term engraftment poor (95,96), and Matrigel is not approved for clinical use (15). In that vein, alternative substrates for hPSC culture, Laminin 521 and vitronectin, are not readily available as pure 3D hydrogels. Laminin 521 has recently been conjugated with silk (Biosilk, Biolamina) to form a generic hydrogel for 3D stem cell and organoid culture (97). However, silk is not native to the human ECM, and Biosilk is not licensed for clinical use. Notably, some hPSC differentiation protocols using vitronectin require dissociating cells mid-process and changing the substrate (98); this disrupts any early 3D structures that may form in vitro.

Transplanted hepatocytes commonly occupy microvascular spaces, and the resulting ischaemic injury triggers vascular and immune responses within the local niche, which usually has a deleterious effect on the transplanted cells (96). This limits cell retention, long-term engraftment, treatment efficacy, and the broader use of cell therapy in liver disease. Other cell delivery methods involve injecting cells encapsulated with degradation-resistant matrices, such as alginate-coated spheroids, delivered into the abdomen (99). This is often a temporary measure to bridge patients with liver failure to liver transplantation; in this situation, the encapsulation limits cell engraftment and repopulation. To investigate the potential of Hepatogel in cell therapy, we injected equal quantities of H-iHeps in Hepatogel and H-iHeps in aqueous solution (standard practice) directly into normal livers of immune-compromised mice (NSG^TM^ mice, n=6 each group per time point, 3 time points). Mice were monitored for human albumin in their serum and then culled for liver histology. Three days after intrahepatic injections, the first group of mice were culled and examined. At post-mortem, it was evident that H-iHeps in Hepatogel were retained at the injection site, visible as pale-coloured cell masses within the liver. These were more distinct than H-iHeps delivered in aqueous solution (0.9% Saline); Fig.6(D).

Respectively, macroscopic appearances corresponded to immunofluorescence staining of engrafted cells that were strongly positive for human albumin at the injection sites (cell masses) but not in other regions of the livers; Fig.6(F). Likewise, the number of H-iHeps was significantly higher in liver histology; Fig.6(G). Significantly higher concentrations of human albumin in mouse serum were also observed over time and corroborated the histological findings; Fig. 6(H). Notably, human albumin remained detectable in mice transplanted with H-iHeps and Hepatogel at Day 6. In contrast, in mice transplanted with H-iHeps in solution, it was undetectable after the second day, suggesting poor cell retention and cell clearance. Nonetheless, beyond 8 days human albumin decreased significantly and was undetectable in both groups. The remaining mice in both groups were culled at 7 days and 2 months after transplantation; no H-iHeps colonies or tumours were seen at the injection sites, demonstrating the clearance of injected H-iHeps by residual innate immunity. Hepatogel was still present.

In summary, Hepatogel improved H-iHeps retention compared to standard therapy when injected directly into the liver. However, sustained H-iHep proliferation and long-term engraftment were not observed. This may have been caused by the immunogenicity of hPSC cells (100), limitations of current iHep technology (101) or immunological responses in recipient mice; NSG^TM^ mice have preserved neutrophil and monocyte function. Limited repopulation of transplanted iHeps remains a universal challenge, and our long-term observations are similar to those of other studies reported in the literature (96). Nonetheless, as biotechnology advances, the derivation of high-quality non-immunogenic lab-derived hepatocytes will improve, and clinically applicable hydrogels will serve as valuable adjuncts in hepatocyte derivation and cell therapy by improving cell delivery, engraftment, and treatment efficacy (21). For clinical translation, a crucial step to improve long-term survival of transplanted cells is to avoid post-transplant immune-mediated reactions. This can be achieved by: (1) using autologous cells, (2) creating a hPSC bank where donors have HLA groups similar to the majority of the population (102), (3) using cell delivery with a supportive matrix (103–105), and (4) using immune suppression after transplantation. The latter is least preferred because lifelong immune suppression in patients can cause long-term side effects such as diabetes, nerve damage, etc. (106); this option may be reserved for animal experimentation. Therefore, strategies that avoid acute or chronic cellular rejection of donor cells in patients should be prioritised.

## CONCLUSION

We have demonstrated proof of concept that bioengineered organ-specific hydrogels can improve the phenotypes of hPSC-derived end-target cells compared with generic hydrogels. Our clinically defined, liver-specific hydrogel (Hepatogel) increased gene expression and metabolic function in induced hepatocytes compared with Matrigel. Being biodegradable, injectable, and composed of clinical-grade biomaterials, Hepatogel may serve as a useful clinical adjunct to improve hepatocyte retention and engraftment for cell therapy.

## FUTURE PERSPECTIVES

We observed that organ-specific hydrogels can produce better end-target phenotypes than Matrigel and its derivatives, a generic mouse sarcoma-derived substrate. Since the benefits of fibrin-only matrices are limited in hPSC culture, several groups have recently attempted to create ECM-enriched fibrin hydrogels for regenerative medicine and cell therapy. Broguiere et al. (23) attempted to mimic Matrigel by enriching fibrin gels with laminin 111 and collagen IV; they subsequently demonstrated that these gels could support the growth of adult stem cells (ASCs) by propagating cystic gastrointestinal organoids (intestinal, pancreatic, and hepatic). Our experiments found that Laminin 111 caused spontaneous differentiation of hPSCs within 1-2 weeks (Supplementary Figure 14), and several groups have independently concluded that this spontaneous differentiation of hPSCs is likely caused by downstream signalling associated with the β1 chain binding to the α3β1-integrin receptor (25–27). Therefore, hydrogels highly enriched in Laminin 111 may be used for ASCs, but they are likely to cause phenotypic instability in hPSCs. Since hPSCs have the potential to generate any cell type in the human body, we believe hPSCs are critical in future efforts to bioengineer any significant tissue mass on a large scale. Hence, clinically translatable biomaterials and culture systems that support stable hPSC expansion and 3D-directed differentiation are cornerstones of regenerative therapies. Synthetic or naturally occurring polymers can be exploited to achieve this. Still, clinical requirements remain significant challenges to overcome – definability, non-toxic polymerisation and gelation, biodegradability for remodelling, biosafety, and ease of use for clinical therapy. Nonetheless, the field’s future is likely to shift closer to biomimicry, using organic-specific and recombinant human matrices to develop biomaterials that yield near-physiological end-target phenotypes. Synthetic alternatives involving complex chemical processes may prove more challenging to translate. In addition, 3D printing technology and microfluidics will play important roles in spatially arranging these biomaterials and growth factors around naïve hPSCs in vitro to recapitulate the physiological niches and microarchitecture, laying the foundation for complete organogenesis.

## Supporting information

Manuscript in MS Word

## ACKNOWLEDGEMENTS

JO was supported by the University of Cambridge (UoC) WD Armstrong graduate studentship, the UoC Engineering in Clinical Practice grant, the National University of Singapore (NUS) Talent Development grant, and the NUS Postdoctoral Development grant. JZ was supported by Trinity College and the Cambridge Commonwealth, European, and International Trust (UoC). Financial support for FC has come from the Engineering and Physical Sciences Research Council (EPSRC, EP/R511675/1); AWJ was supported by the Isaac Newton Trust (to AEM), the Rosetrees Trust (M787 to AEM and AWJ) and the Wellcome Trust Institutional Translational Partnership Award 222062/Z/20/Z (to AEM, AWJ and SS). SS was funded by the British Heart Foundation FS/18/46/33663. We also acknowledge support from the Wellcome Trust (DRP 226795/Z/22/Z; AEM and SS). We gratefully acknowledge Dr MA Al Fahad for assistance with generating the heatmap plot included in this manuscript.

## Ethics approval statement

This study did not involve human subjects. The National University of Singapore Institutional Animal Care and Use Committee (IACUC) granted ethical approval for animal research, protocol number R20-0822.

