## Supplementary material for "A clinically defined and xeno-free hydrogel system for regenerative medicine": Manuscript in MS Word

*Stemness Index (SI) =*

*Total number of Stem Cells Identified*

*Total number of Cells In Population*

The SI values range from 0 to 1, and the equation was incorporated into the pipeline. Positive hits were checked using brightfield microscopy.

Table 1: List of primary antibodies.

| ***Primary Antibodies***  ***(species raised in)*** | ***Vendor*** | ***Catalogue no.*** | ***Dilution factor*** |
| --- | --- | --- | --- |
| OCT4 (rabbit) | Thermo Fisher | 710788 | 1:250 |
| NANOG (rabbit) | Abcam | ab109250 | 1:100 |
| SOX2 (rabbit) | Thermo Fisher | PA1-094 | 1:100 |
| SOX2 (goat) | RnD Systems | AF2018 | 10 μg/ml |
| Albumin (goat) | Bethyl | A80-129A | 1:100 |
| CD147 (mouse) | Abcam | Ab666 | 1:100 |
| HNF1A (rabbit) | Cell Signalling | D7Z2Q | 1:400 |
| HNF4A (rabbit) | Thermo Fisher | MA5-14891 | 1:2000 |
| Cyp2A6 (chicken) | University of Eastern Finland | Collaborator | 1:200 |
| E-cadherin (mouse) | Santa Cruz | SC-31021 | 1:100 |
| Troponin-T (goat) | Abcam | ab64623 | 1:100 |
| Smooth Muscle α-Actin (mouse) | Dako | M0851 | 1:100 |
| Nestin (mouse) | Sigma | N5413-100UG | 1:500 |
| Β-catenin (rabbit) | Thermo Fisher | 14-2567-82 | 1:250 |

Table 2: List of secondary antibodies

| ***Secondary Antibodies*** | ***Vendor*** | ***Catalogue no.*** |
| --- | --- | --- |
| AlexaFluor 488 donkey anti-goat | Thermo Fisher | A11055 |
| AlexaFluor 488 donkey anti-rabbit | Abcam | ab150073 |
| AlexaFlour 488 donkey anti-rabbit | Thermo Fisher | A21206 |
| AlexaFluor 564 donkey anti-mouse | Thermo Fisher | A10037 |
| AlexaFluor 647 donkey anti-rabbit | Abcam | ab150075 |
| AlexaFluor 647 donkey anti-mouse | Thermo Fisher | **A32795** |
| AlexaFluor 647 donkey anti-mouse | Thermo Fisher | **A31571** |

Table 5.3: Sequences for forward and reverse primers used.

| ***Target gene*** | ***Forward primer sequence*** | ***Reverse primer sequence*** |
| --- | --- | --- |
| ABCC2 | TTCGTTCCAGACGCAGTCCAGG | CCAGGAGCCATAGGTAGCCCAA |
| APOB | TGTTAGGACACCAGCCCTCCA | GCCCGAAGGCTGAAATGGTCT |
| AFP | AACTATTGGCCTGTGGCGAGG | TGAAGCATGGCCTCCTGTTGG |
| BAAT | TGTGCCTCAACGACCCACGAT | GGGGGAGACATTCCGCCATGAA |
| CEBPA | AAGTCGGTGGACAAGAACAGCAA | TTGTCACTGGTCAGCTCCAGCA |
| CYP27A1 | GCTGTTCGTTCAAGGCTATGCCC | ACTGGGTACTTGCCCTCTTGCC |
| CYP3A4 | TGTGCCTGAGAACACCAGAG | GTGGTGGAAATAGTCCCGTG |
| CYP3A5 | ACAAGACCCCTTTGTGGAGAGC | TTGTTTGTCGTTGAGGCGACTT |
| CYP3A7 | TGGACCCAGAAACTGCATTGGC | ATCAGGCTCCACTTACGGTCTCA |
| CK7 | AAGCAGGATATGGCACGGCAG | CACTCCATCTCCAGCCAACCG |
| CK19 | GGCGATGTGCGAGCTGATAGT | CGGTAGGTGGCAATCTCCTGC |
| FOXA1 | CCTTCAACCACCCGTTCTCCA | CGGGCAACGTAGAGCCGTAAG |
| FOXA2 | GCCTACGAACAGGTGATGCACTA | CAGGCCCGTTTTGTTCGTGAC |
| GATA6 | AGCACCAATCCCGAGAACAGC | GTCGCACGGAGGACGTGACTT |
| GATA4 | TCCATCCACCCTGTCCTCTCG | GCTTGGAGCTGGTCTGTGGAG |
| G6PC | TACAGCAACACTTCCGTGCCCC | GCACAGCCCAGAATCCCAACCA |
| GGT | ACAACCAGCTTCTGCCCAACG | GACGCGATCTGGGTGTGATGG |
| HNF4A | GAGATGCTGCTGGGAGGGTCC | GGGTCTCAGGGGTGGACATCT |
| LGR5 | CCTGCTTGATGGCTGGCTGAT | CACTGCTGCGATGACCCCAAT |
| PDK4 | TACAGCAACACTTCCGTGCCCC | GCACAGCCCAGAATCCCAACCA |
| SOX9 | CAACAGATCGCCTACAGCCCC | CGTACTGTGAGCGGGTGATGG |
| TAT | TGGAAACCTGCCTACAGACC | TAGCTTCTAGGGGTGCCTCA |
| TF | CCGATGGTCCCAGTGTTGCTT | AACTCTGCCACCACAGGCTTC |
| TBX3 | GGGACACTGGAAATGGCCGAA | GGAGAAGAAGCCTGGGCGAAG |
| MUC2 | GCTGACGAGTGGTTGGTGAATG | GATGAGGTGGCAGACAGGAGAC |

***2.14 Enzyme-linked immunosorbent assays (ELISA)***

Laminin concentrations in supernatants were measured using a laminin ELISA kit (Insight Biotechnology, #Boster Pikokine^TM^ EK0434). Acellular FLGs and fibrin gels were synthesised, and sterile DMEM/F12 media was added 30 min after gelation. Media was collected the next day (Day 1) and every other day for two weeks. Supernatants were stored at -20 ^o^C until they were analysed alongside stock concentrations of laminin for comparison using the Laminin ELISA kit per the manufacturer's instructions. Sample absorbance readings were then taken using a FLUOstar® Omega plate reader (BMG Labtech Ltd, UK) at 450 nm. The following equation estimated the residual amount of laminin in FLGs:

$$1 - \frac{Total Laminin 521 in supernatant after hydrogel gelation (weight)}{Measured Laminin 521 in stock solution added to the hydrogel (weight)}$$

Albumin concentrations in supernatants were measured using a human albumin ELISA quantification kit (Bethyl Lab, #E80-129). Sample absorbance was measured with a Synergy HTX microplate reader (BioTek, USA). Supernatants from primary human hepatocytes (PHH) were collected 24 hr after the completion of cell recovery. Supernatants from iHeps were collected 2 days after (Day 22) the end of the differentiation protocol (Day 20); media were refreshed every other day as per the differentiation protocol.
