## Supplementary Data for "A clinically defined and xeno-free hydrogel system for regenerative medicine"

### Supporting Data

#### Dissolution of cellularised Laminin 521-enriched fibrin hydrogels (Alphagel)

Accutase (Sigma, #A6964-100ML), Cell Dissociation Buffer (ThermoFisher, #13151014), and TrypLE Express (Gibco, #12604013) were added directly to samples as per manufacturers' recommendations. EDTA (ThermoFisher, #17892) was dissolved in sterile PBS (Sigma, #806552) to a concentration of 0.5 mM, then syringe-filtered and stored at 4°C before use [1]. Sterile Trypsin (ThermoFisher, #A15090046) was dissolved in PBS (without calcium and magnesium) to a final concentration of 0.25% before use. Nattokinase (Simply Pure, #SKUX000HJG6BT) was dissolved in PBS to desired concentrations, syringe-filtered, then stored at 4°C before use. All dissolution reagents were added at a volume of 1ml per well in a 24-well plate.

Nattokinase required the least amount of time to dissolve fibrin-laminin hydrogels, followed by Trypsin, TrypLE Express and Accutase, respectively; Table S1. However, when tested with HPSCs, cells exposed to Nattokinase failed to proliferate when re-seeded back into 3D fibrin-laminin hydrogels. Therefore to investigate if Nattokinase adversely affected HPSCs, several concentrations of Nattokinase were added to HPSCs cultured in standard conditions (2D Geltrex-coated tissue culture plates).

| Reagent | Acellular fibrin-laminin dome (50µl) dissolution times |
|---|---|
| Accutase | 30 minutes - 35 minutes |
| Cell Dissociation Buffer | No effect after 60 minutes |
| EDTA | No effect after 60 minutes |
| Trypsin | 15 minutes - 20 minutes |
| TrypLE Express | 20 minutes - 25 minutes |
| Nattokinase: 1000 units/ml | 10 minutes - 15 minutes |
| Nattokinase: 750 units/ml | 15 minutes - 20 minutes |
| Nattokinase: 500 units/ml | 15 minutes - 20 minutes |

**Table S1**: Time required to dissolve 50µl FLG domes with various reagents.

Interestingly, changes in cell morphology were observed within 10-15 minutes of exposure to Nattokinase (Figure S1). Cells appeared unhealthy and began to detach. Trypan Blue cell counts showed that HPSCs exposed to Nattokinase exhibited low viability, whereas cells unexposed to Nattokinase maintained high viability (> 95%). The decrease in cell viability was directly related to increasing Nattokinase concentrations: 1000 U/ml (0-10% cell viability), 750U/ml (10-20% cell viability), and 500 U/ml (10-20% cell viability).

Since Nattokinase was found to be cytotoxic to HPSCs, Trypsin and TrypLE Express were subsequently evaluated for their effects on cell viability. This revealed that although TrypLE Express required longer to dissolve fibrin-laminin hydrogels, it resulted in higher cell viability compared with Trypsin (90-95% vs 70-80%, respectively). Accutase had similar cell viability with TrypLE Express but required significantly longer dissolution times. Therefore, TrypLE Express was determined to be the best dissolution agent for Laminin 521-enriched fibrin hydrogels.

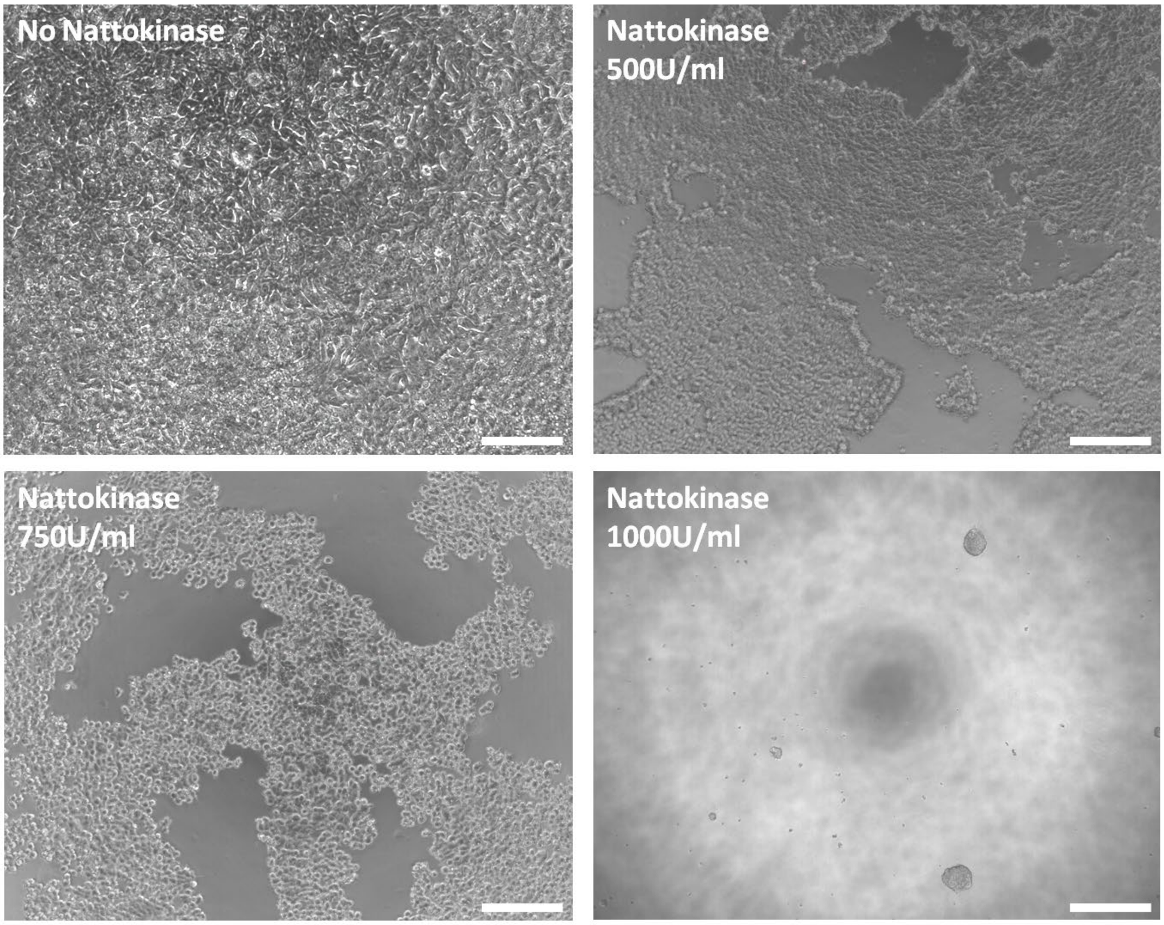


**Figure S1**: Effects of Nattokinase on fully confluent human embryonic stem cells on Geltrex. Scale bar = 100 μm.
