## Supplementary Figures for "A clinically defined and xeno-free hydrogel system for regenerative medicine"

### Supplementary Figure 1

A

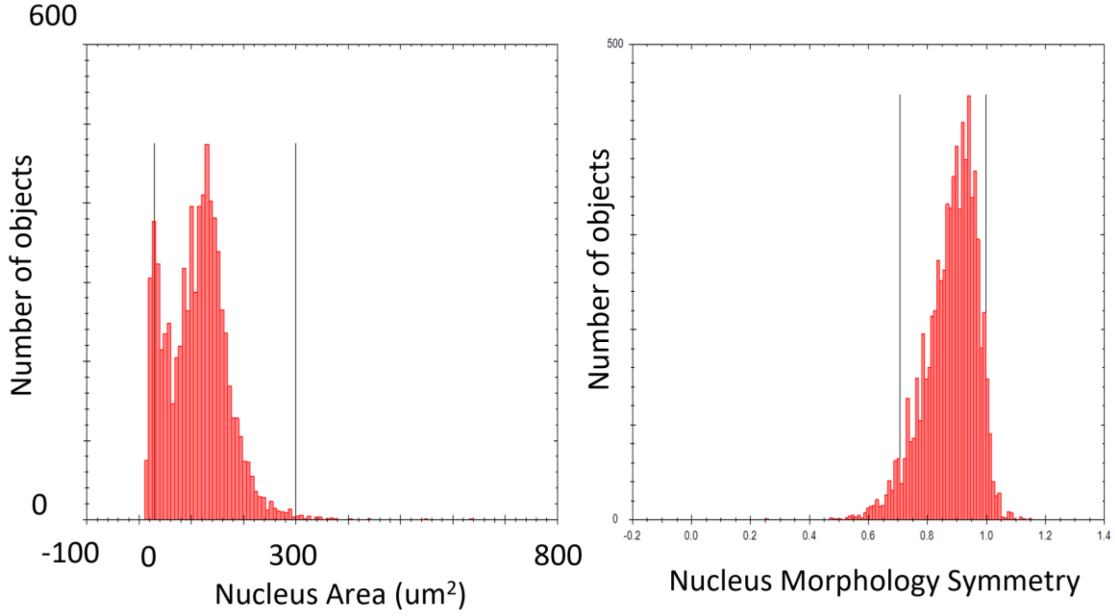


B

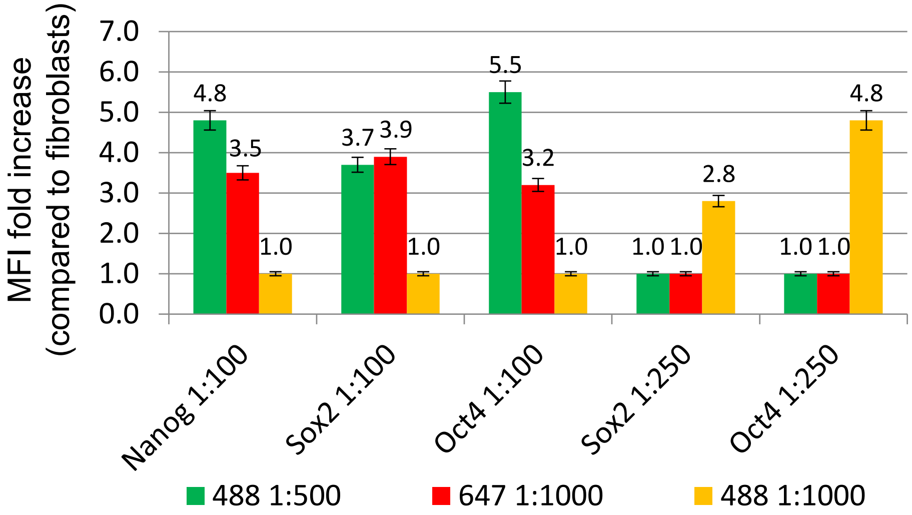


Supplementary Figure 1. A) Histograms were used to characterise the variation in hPSC nuclear area (left) and nuclear roundness (right). A Receiver Operating Characteristic (ROC) curve was then plotted using the MFIs of OCT4, NANOG, and SOX2 in hPSCs and fibroblasts to evaluate the discriminatory ability of the stem cell markers. ROC curve analyses of hPSC nuclear properties alone were found to be poorly discriminatory between hPSC and fibroblasts. The Youden Index was calculated for each stem cell marker to determine optimal MFI cut-offs (below) for positively identifying hPSCs. “Building blocks” in Harmony (Perkin Elmer, USA) were then sequenced to first identify single cells based on the nuclear properties illustrated above, and to exclude clumps and artefacts. Then, from this population of cells, hPSC identification was performed using the MFI cut-offs of stem cell markers i.e. OCT 4 > 3221, NANOG > 3536, and SOX2 >1977. The cells identified by this pipeline were marked as hPSCs. The Stemness Index was then calculated by dividing the number of hPSCs identified by morphology and pluripotency markers by the total number of cells identified by DAPI. This was validated by using the pipeline to identify hPSCs, which were first counted manually, then plated with varying numbers of fibroblasts, which were also counted manually (using Tryptophan Blue). B) Fold increase in mean fluorescent intensity (MFI) in hPSCs cells compared to fibroblast at various primary and secondary antibody concentrations. At specific antibody concentrations, the fold increase in MFI clearly discriminated pluripotent hPSCs from human fibroblasts.

### Supplementary Figure 2

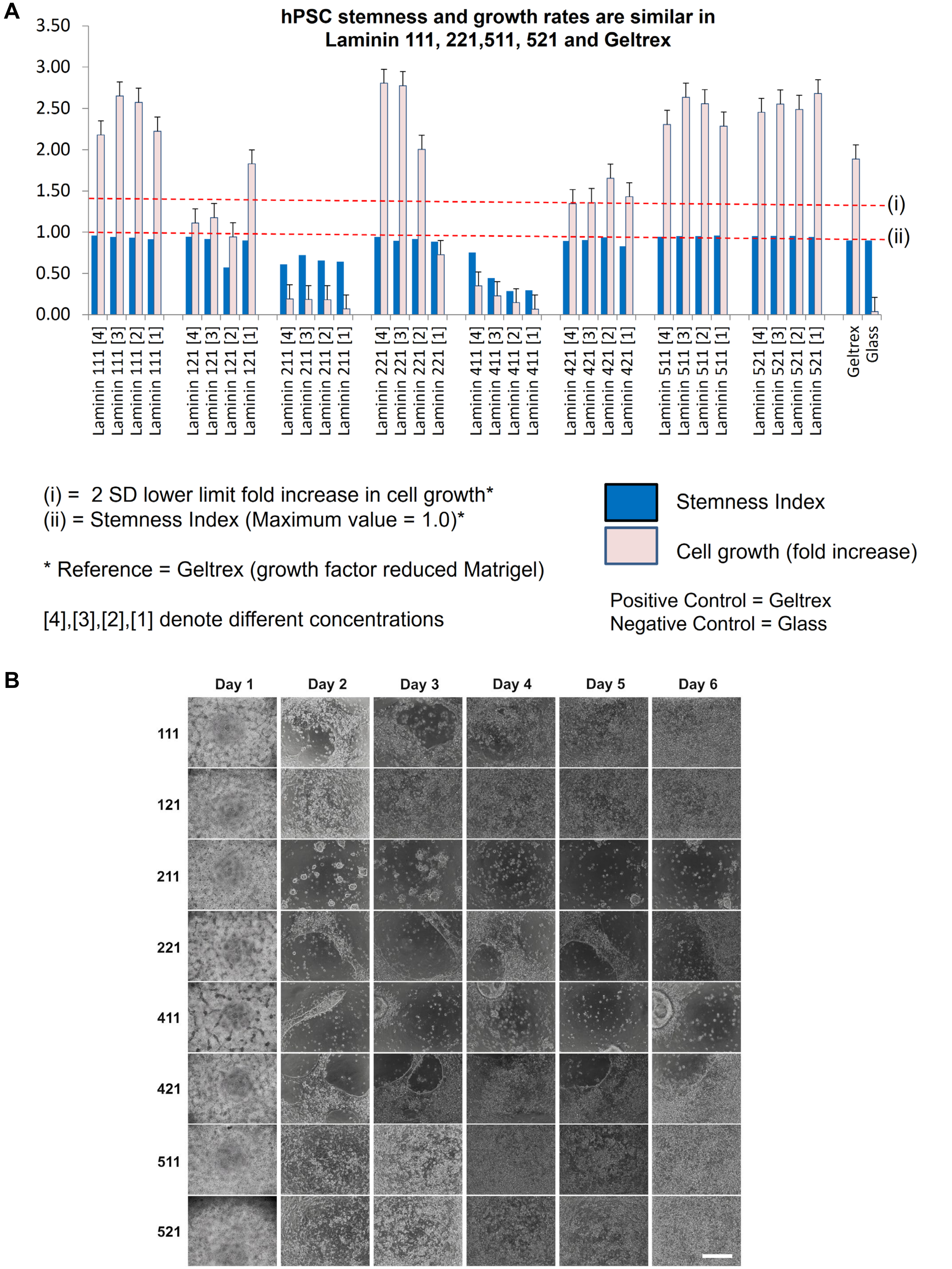


Supplementary Figure 2. A) An example of a focused screening panel. Calculated stemness index (SI) and growth rates of hPSCs on laminins after 1 week in culture (*n* = 3 biological samples with 3 experimental replicates at each concentration. B) Bright-field microscopy of hPSCs on various laminin isoforms (y-axis) at 0.75 µg/ml over time (x-axis) validating results from A). Scale bar = 100 µm.

### Supplementary Figure 3

A

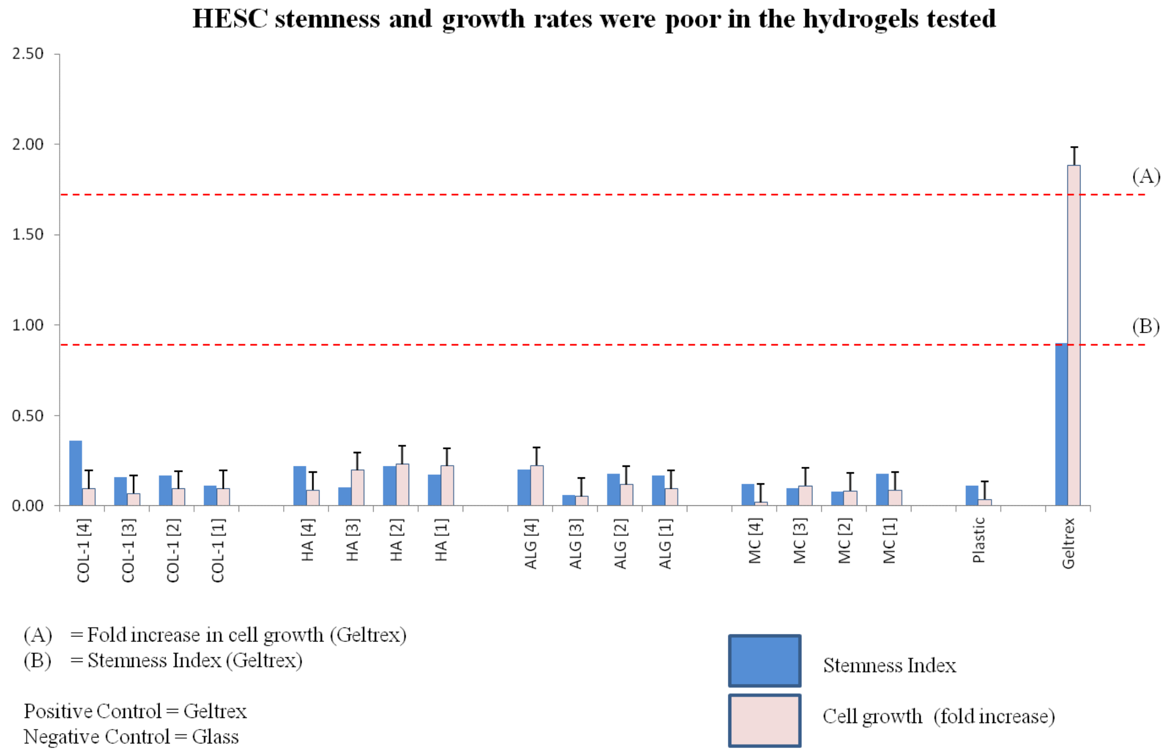


B

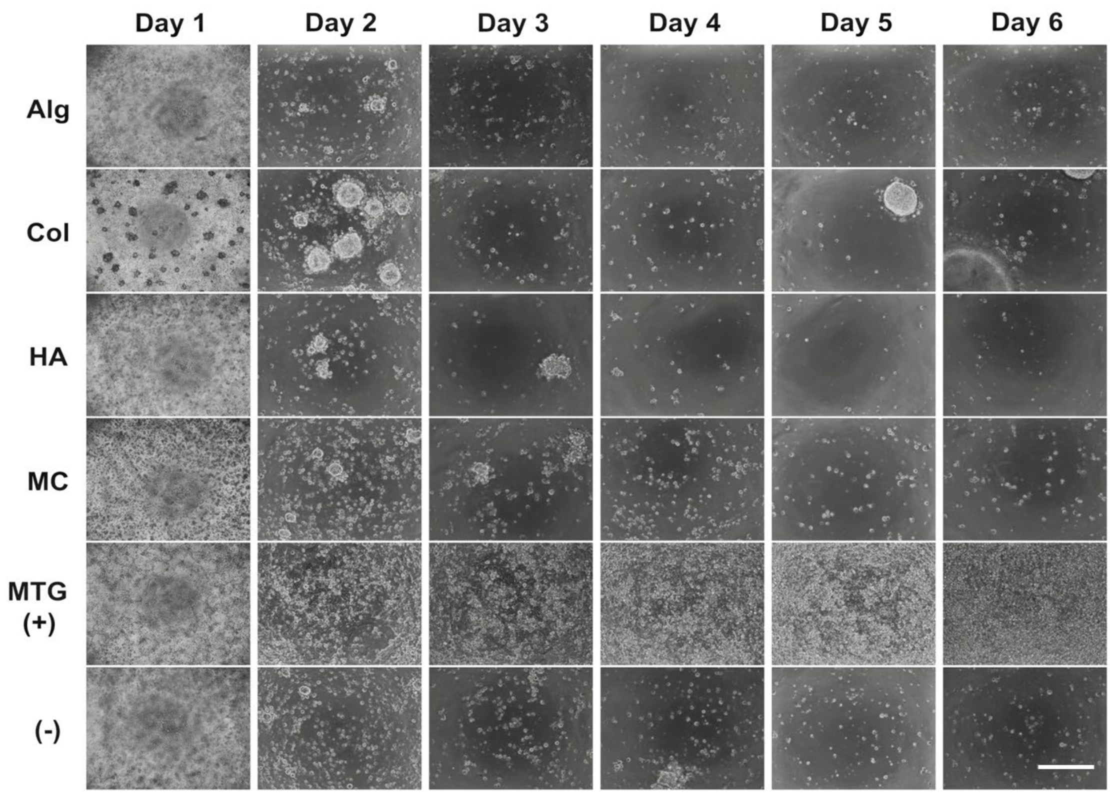


Supplementary Figure 3. A) An example of a focused screening panel of various polymeric proteins. Calculated stemness index (SI) and growth rates of hPSCs on laminins after 1 week in culture ($n$ = 3 biological samples with 3 experimental replicates at each concentration. B) Bright-field microscopy of hPSCs on various polymeric proteins (y-axis) over time (x-axis) validating results from (A). Alg = alginate, Col = collagen-1, HA = hyaluronic acid, MC = methylcellulose, MTG (+) = Geltrex (positive control), and (-) = Tissue culture treated glass bottom plate (negative control). Scale bar = 100 μm.

### Supplementary Figure 4

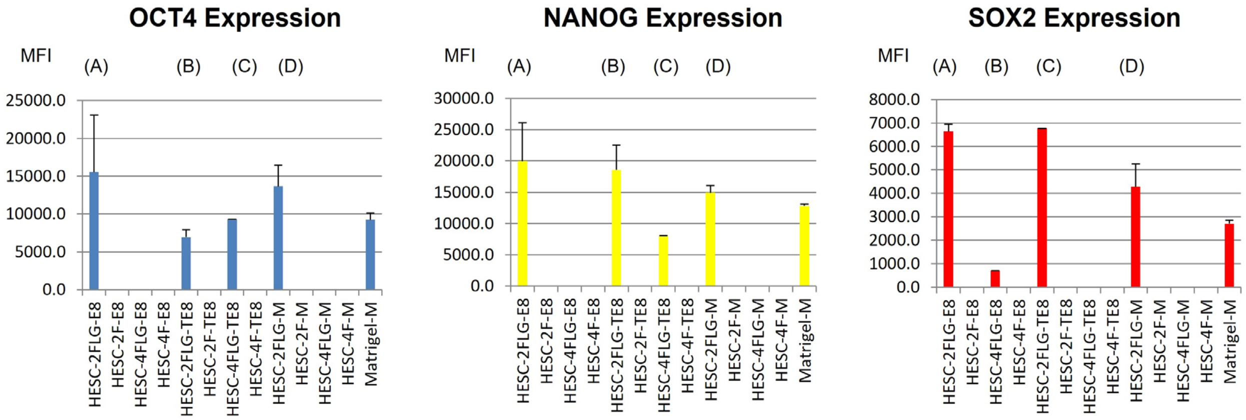


| | |
|---|---|
| (A) vs Matrigel | p = 0.27 |
| (B) vs Matrigel | p = 0.13 |
| (C) vs Matrigel | Not applicable |
| (D) vs Matrigel | p = 0.13 |
| Stem. Index (A) | 0.62 |
| Stem. Index (B) | 0.59 |
| Stem. Index (C) | 0.87 |
| Stem. Index (D) | 1.0 |
| Stem. Index (MTG) | 0.98 |

| | |
|---|---|
| (A) vs Matrigel | p = 0.13 |
| (B) vs Matrigel | p = 0.13 |
| (C) vs Matrigel | Not applicable |
| (D) vs Matrigel | p = 0.13 |
| Stem. Index (A) | 0.98 |
| Stem. Index (B) | 1.00 |
| Stem. Index (C) | 0.54 |
| Stem. Index (D) | 1.00 |
| Stem. Index (MTG) | 1.00 |

| | |
|---|---|
| (A) vs Matrigel | p = 0.13 |
| (B) vs Matrigel | Not applicable |
| (C) vs Matrigel | Not applicable |
| (D) vs Matrigel | p = 0.13 |
| Stem. Index (A) | 0.79 |
| Stem. Index (B) | 0.01 |
| Stem. Index (C) | 0.50 |
| Stem. Index (D) | 0.94 |
| Stem. Index (MTG) | 0.72 |

Supplementary Figure 4: Stem cell marker expression (MFI) presented as mean ± standard error of the mean (S.E) and stemness index (SI) in various culture conditions after 3-4 months. Bar chart x-axis label key: (i) cell type, (ii) fibrin concentration (2 mg/ml or 4 mg/ml), (iii) gel-type = fibrin-laminin hydrogel (FLG) or fibrin only (F), and (iv) media type = mTeSRs/TeSR2 (M), TeSR-E8 (TE8), and E8 (E8). 2FLG is representative of Alphagel. HESCs cultured in 2 mg/ml fibrin gel with FLG in E8 media. The absence of bars and/or S.E. was because cells died or differentiated in other repeats and no longer expressed stem cell markers. p values derived using the Mann Whitney U test (MFI), Chi-square test (SI) or Fisher test (when $n < 5$ for any cell).

### Supplementary Figure 5

| | Corning Matrigel (hESC qualified) from Fisher Scientific | Alphagel using **clinical grade** recombinant human Laminin 521 from Biolamina | Alphagel using **research-grade** recombinant human Laminin 521 from Biolamina |
|---|---|---|---|
| Cost of matrix on Cambridge EDRS | £500.50 for 10 ml (ID: 10365602) | £546.26 (ID: CT521-0501) | £364.17 (ID: LN521-05) |
| (a) Cost per microlitre | 5 p | 5.5 p | 3.64 p |
| (b) Matrix volume per 50 µl dome | 33.3 µl | 25 µl | 25 µl |
| (c) Matrix cost per 50 µl dome (a) x (b) | £1.67 | £1.37 | £0.91 |
| (d) Cost of human thrombin (50 µl dome) | NA | 10.4 p (£78.84 for 1 gm) | 10.4 p (£78.84 for 1 gm) |
| (e) Cost of human fibrinogen (50 µl dome) | NA | 1.7 p (£171.00 for 1 gm) | 1.7 p (£171.00 for 1 gm) |
| (f) Total of 50 µl dome: (c)+(d)+(e) | £1.67 | £1.49 | £1.03 |
| Cost per ml of gel: (f) x 20 | £33.40 | £29.80 | £20.60 |
| Potential cost savings (vs Matrigel) | - | **10.8%** | **38.3%** |

Supplementary Figure 5: Cost-analysis to synthesize a 50 µl hydrogel dome for 3D cell culture; Matrigel vs Alphagel with clinical grade laminin vs Alphagel with research grade laminin.

### Supplementary Figure 6

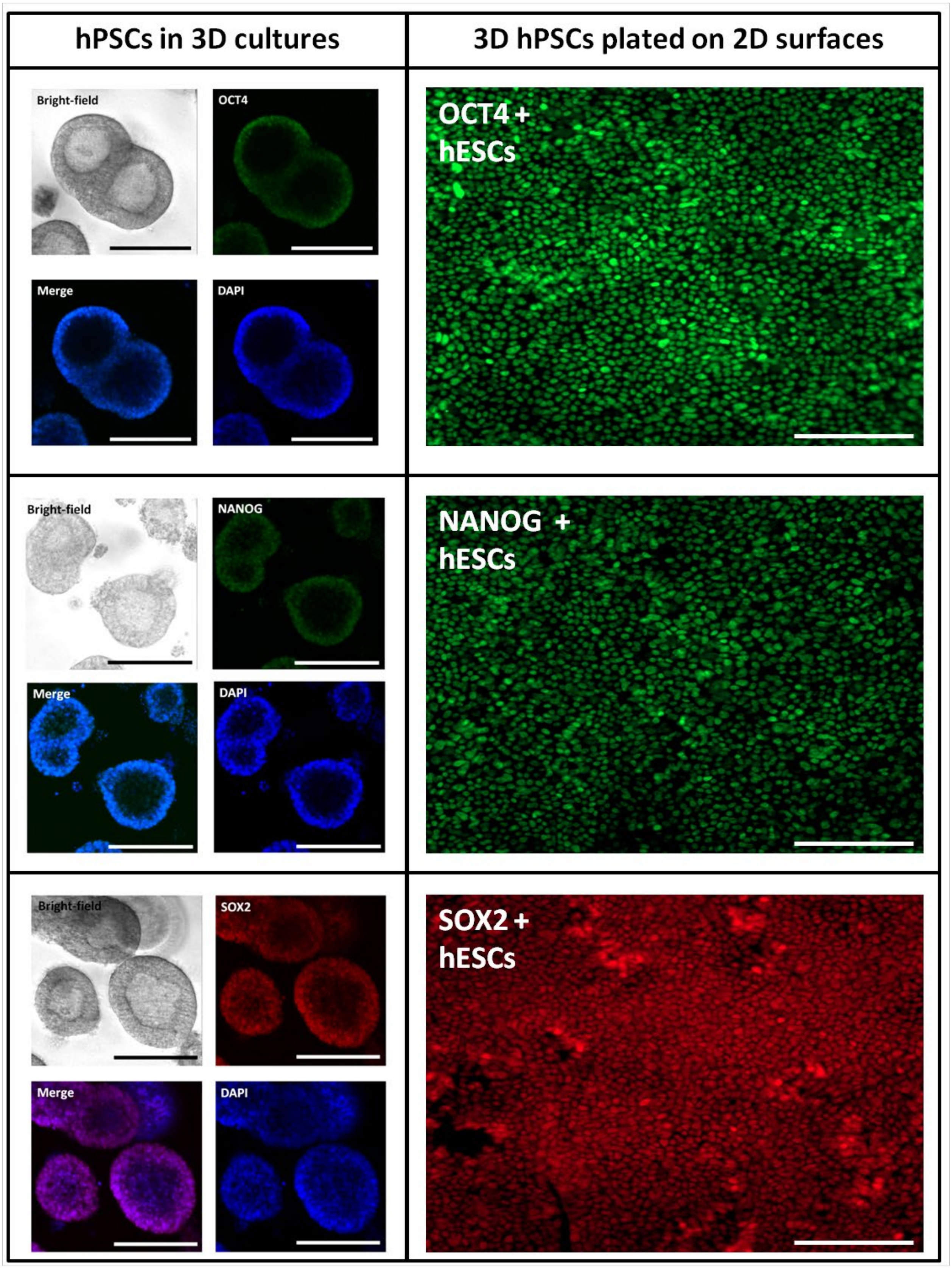


Supplementary Figure 6: Left column: Bright-field and immunofluorescent images with positive nuclear staining of hPSC in 3D. Right column: hPSCs derived from 3D cell aggregates and cultured as a 2D monolayer. Scale bar = 200 μm. Cells continue to proliferate and express nuclear markers of pluripotency.

#### Supplementary Figure 7

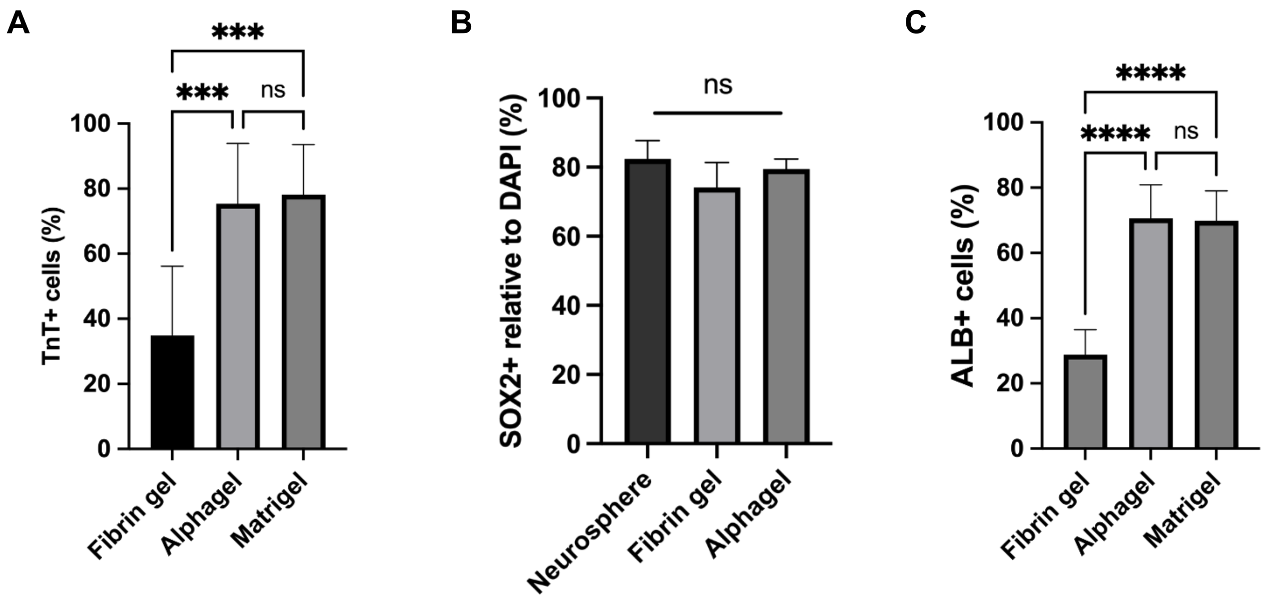


Supplementary Figure 7: A) Percentage of Troponin T-positive (TnT+) cells after 14 days of cardiac differentiation. B) Percentage of SOX2 positive (SOX2+) neural progenitor cells after 16 days of neural differentiation, and C) percentage of albumin positive (ALB+) cells after 22 days of hepatic differentiation.

### Supplementary Figure 8

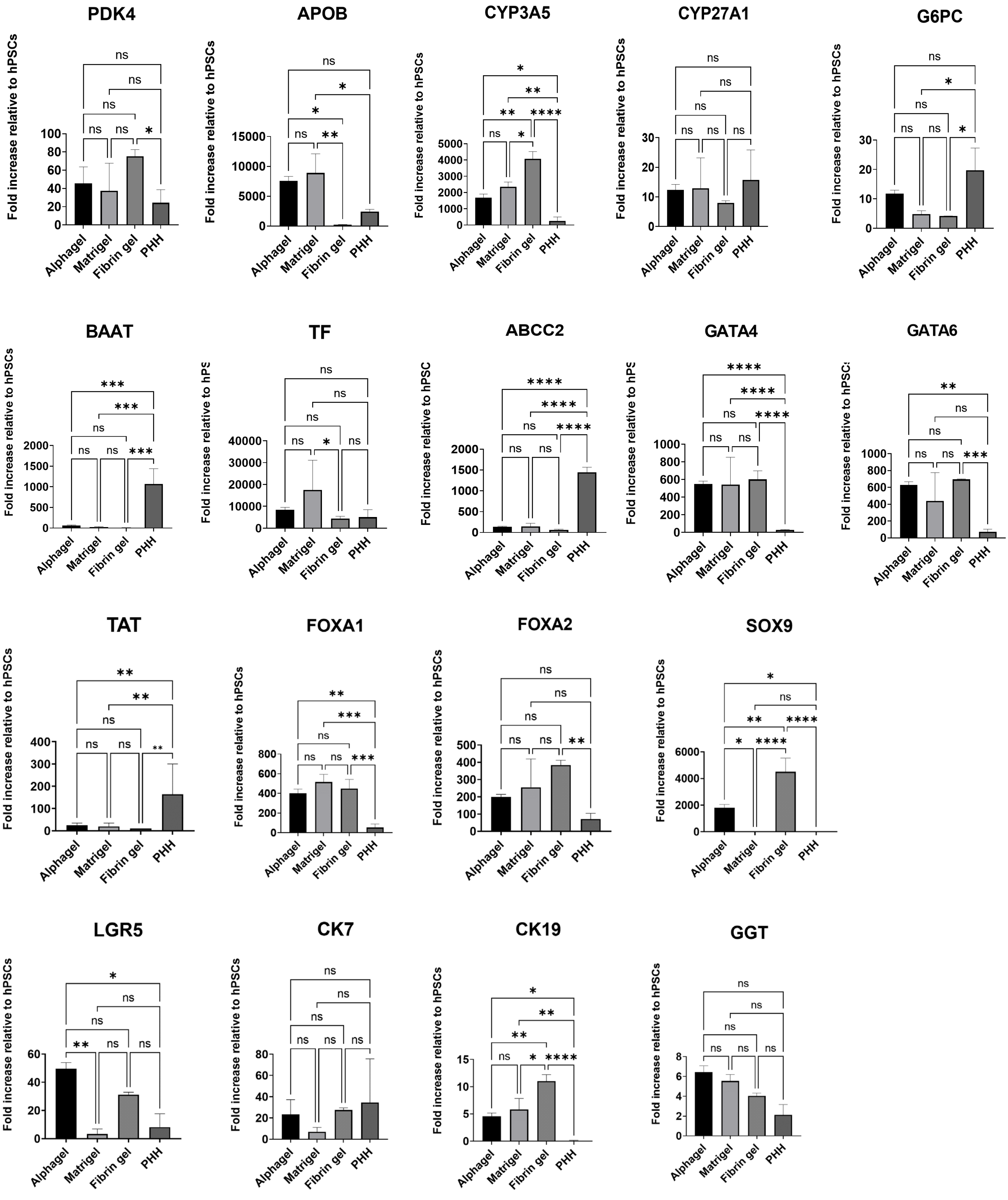


Supplementary Figure 8: Gene expression data of the remaining 18 of 24 liver-associated genes analysed.

### Supplementary Figure 9

| Microscopic lesions | Score |
|---|---|
| 1. Infiltrate, inflammatory cell, Polymorphs | |
| 2. Infiltrate, inflammatory cell, Mononuclear cells | |
| 3. Haemorrhage | |
| 4. Suppuration | |
| 5. Fibrosis | |
| 6. Myofibers degeneration/necrosis | |
| 7. Adipose tissue necrosis | |
| 8. Myofibers regeneration* | |
| 9. Granulation tissue* | |
| Total Score | |

Supplementary Figure 9: Histology scoring tool used to assess inflammation on histopathological evaluation. Each feature of inflammation is scored on an ordinal scale from 0 to 5 (0 = no abnormality, 1 = minimal abnormality, 2 = mild abnormality, 3 = moderate abnormality, 4 = severe abnormality, and 5 = marked abnormality). All nine components are then summated to provide the total score which correlates directly to the severity of inflammation.

### Supplementary Figure 10

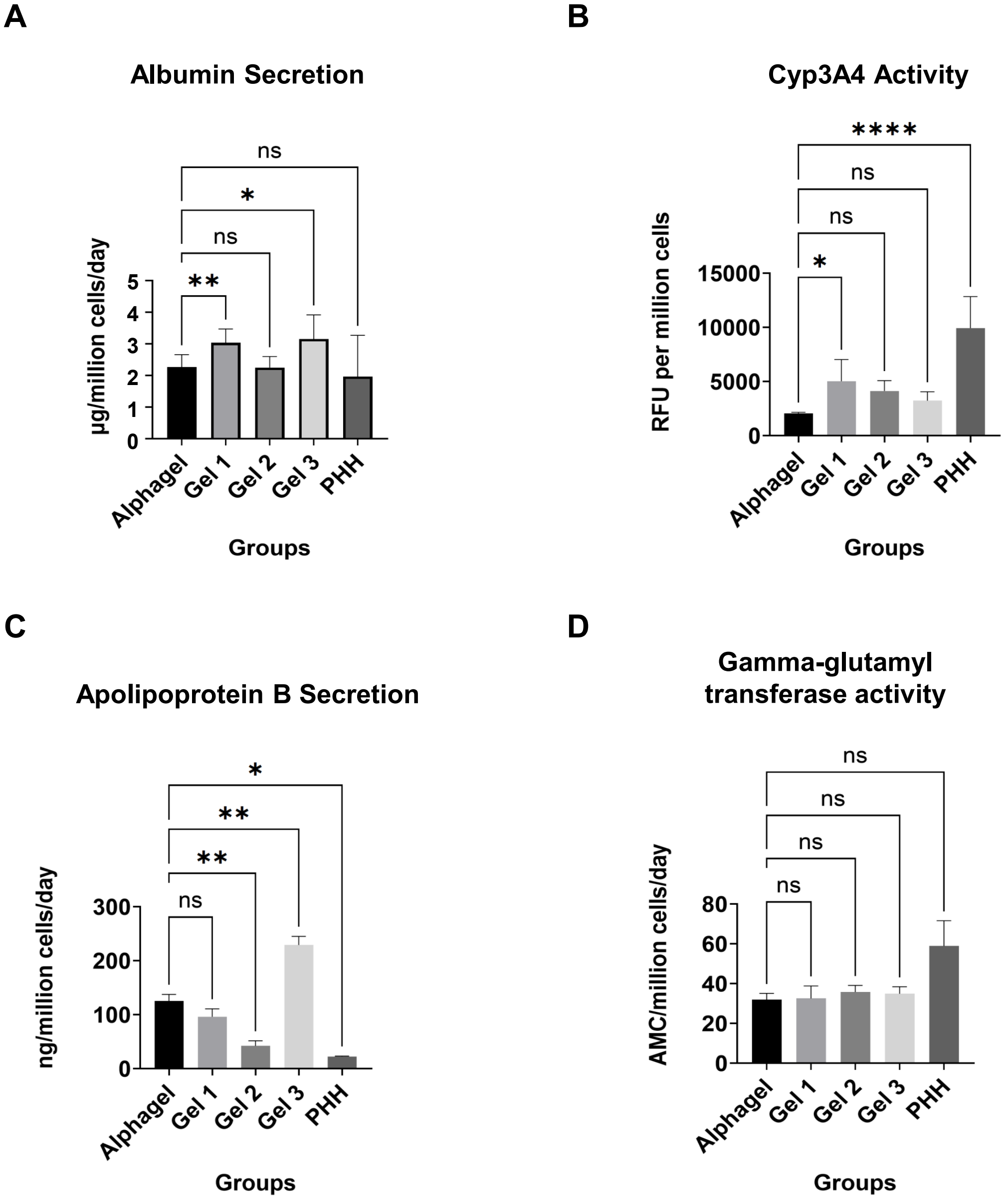


Supplementary Figure 10: Functional assays of i-Heps cultured in hydrogels containing different ratios of Laminin 521: Laminin 111: Laminin 411, respectively. A) Albumin secretion and B) Cyp3A4 activity were highest in Hepatogel (Gel 1, 5:2:1). C) Apolipoprotein B was highest in Gel 3 (5:1:2), and no significant differences were noted in gamma-glutamyl transferase activity across all conditions D). The ratios for Gel 2 were 4:2:2.

### Supplementary Figure 11

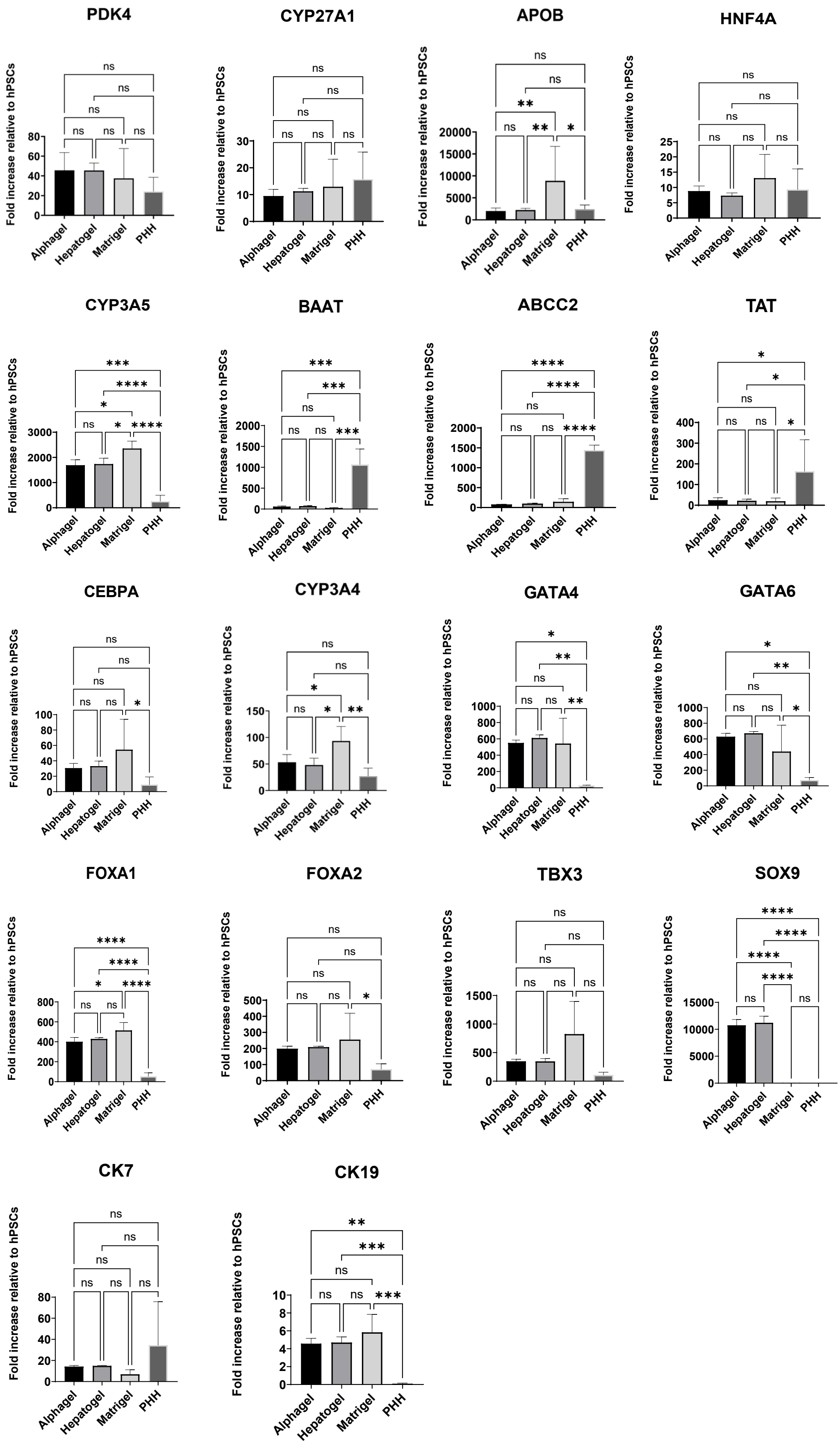


Supplementary Figure 11: Remaining gene panel of iHeps cultured in Hepatogel, Alphagel, and Matrigel vs PHH. iHeps were harvested between Day 22-24 of the hepatic differentiation process.

### Supplementary Figure 12

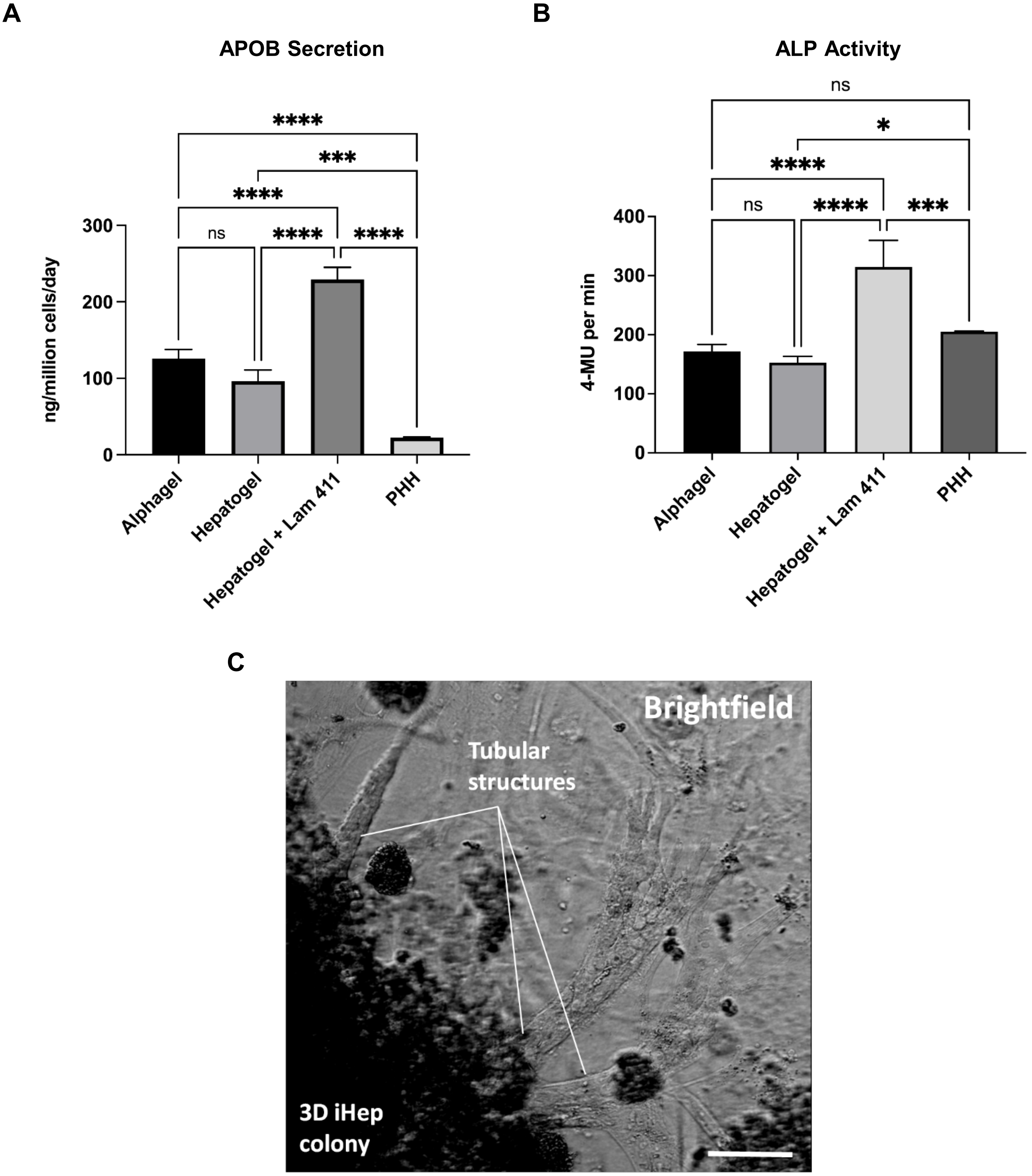


Supplementary Figure 12: A) ELISA of culture medium for Apolipoprotein B (APOB), taken from iHeps 2 days after the end of the hepatic differentiation protocol, and from PHH culture media 2 days after plating. B) Alkaline Phosphatase (ALP) activity measured by colourimetric assay on culture medium taken 2 days the end of the hepatic differentiation protocol. C) Brightfield microscopy of 3D tubular branching strictures observed when Laminin 411 concentrations are increased. Scale bar = 100 μm.

#### Supplementary Figure 13

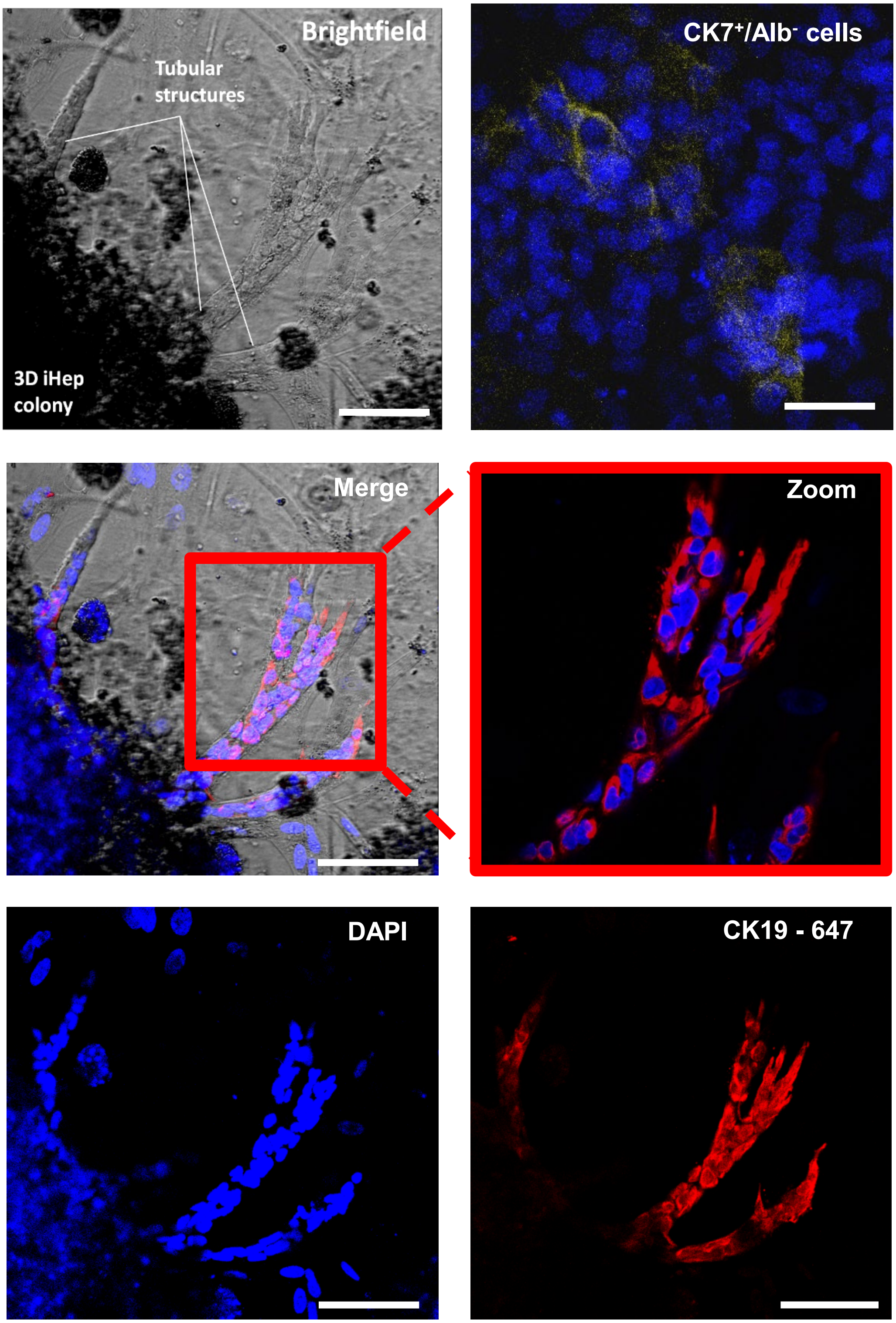


Supplementary Figure 13: Tubular structures comprising of cells that are Cytokeratin 7 (CK7) positive (yellow), Cytokeratin 19 (CK19) positive (red) and albumin negative. DAPI = blue. Scale bar = 100 μm.

### Supplementary Figure 14

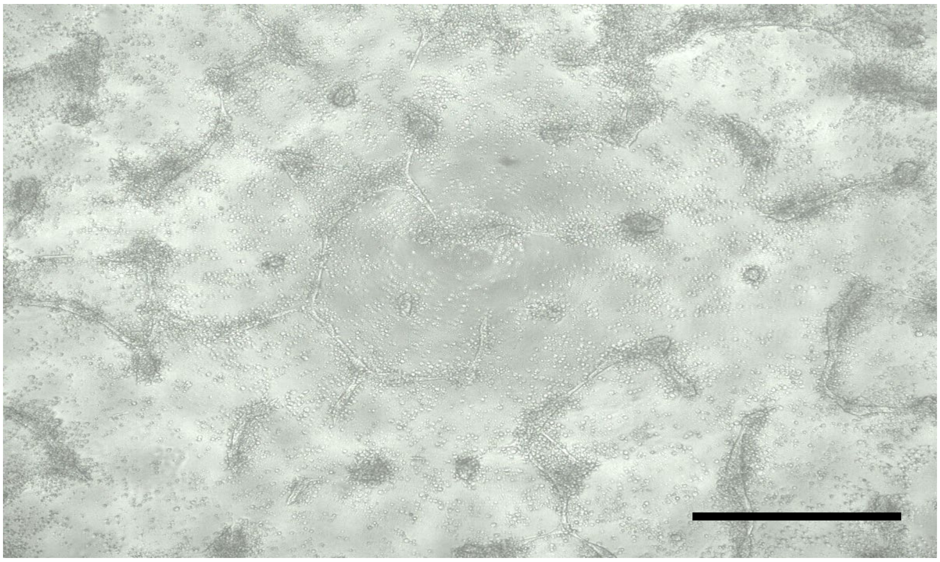

Supplementary Figure 14: Tube-like formations in hESCs on cultured on Laminin 111 with mTESR media on 2nd passage, x40 magnification. Scale bar = 250 μm.
